# Comparative Transcriptomic Analysis of Perfluoroalkyl Substances-Induced Responses of Exponential and Stationary Phase *Escherichia coli*

**DOI:** 10.1101/2025.02.18.638913

**Authors:** Molly E. Wintenberg, Olga B. Vasilyeva, Samuel W. Schaffter

## Abstract

Per- and polyfluoroalkyl substances (PFAS) are highly stable chemical contaminants of emerging concern for human and environmental health due to their non-natural chemistry, widespread use, and environmental persistence. Despite conventional metrology, mitigation strategies, and removal technologies, the complexity of this growing problem necessitates the need for alternative approaches to tackle the immense challenges associated with complex environmental PFAS contamination. Recently, biology has emerged as an alternative approach to detect and mitigate PFAS and understand the molecular-level responses of living organisms, including microorganisms, to these compounds. However, further study is needed to understand how microorganisms in different environments and growth phases respond to PFAS. In this study, we performed RNA sequencing at mid-exponential, early stationary phase, and late stationary phase of bacterial growth to determine the global transcriptional response of a model chassis, *Escherichia coli* MG1655, induced by two PFAS, perfluorooctanoic acid (PFOA) and perfluorododecanoic acid (PFDoA), and equivalent non-fluorinated carboxylic acids (NFCA), octanoic acid and dodecanoic acid. Differential gene expression analysis revealed PFOA and PFDoA induced distinct changes in gene expression throughout cultivation. Specifically, we identified significant changes in expression of the formate regulon and sulfate assimilation at mid-exponential phase and ferrous iron transport, central metabolism, the molecular chaperone network, and motility processes during stationary phase. Importantly, many of these changes are not induced by NFCAs. In summary, we found PFAS induced a system-level change in gene expression, and our results expand the understanding of bacterial-PFAS interactions that could enable the development of future real-time environmental monitoring and mitigation technologies.

**Importance:** The prevalence and persistence of PFAS in the environment is a growing area of concern. However, little is understood of the impacts of PFAS on the environment, particularly impacts on microorganisms that play pivotal roles in nearly every ecosystem. Thus, comprehensive measurements that provide systems-level insight into how microorganisms respond and adapt to PFAS in the environment are paramount. Here, we use RNA sequencing to study the global transcriptional response of *E. coli* MG1655 to two PFAS and non-fluorinated equivalent compounds across growth phases. We find that PFAS induce system-level changes in metabolic, transport, and gene regulatory pathways, providing insight into how these non-natural chemicals interact with a model bacterium. Additionally, the transcriptomic dataset associated with this work provides the community with PFSA-specific gene expression patterns and possible PFAS degradation pathways for the development of future whole-cell biosensors and mitigation efforts.

## INTRODUCTION

Per- and polyfluoroalkyl substances (PFAS) are a class of fluorinated compounds with various structures conferring strong resistance to degradation or decomposition in the environment (1). Widespread industrial production of fire-fighting foam and non-stick coatings and use in other consumer products of pesticides, cleaning materials, and personal care products, has led to substantial release of PFAS into the environment, raising concerns about their environmental persistence and bioaccumulation (2, 3). PFAS have been detected ubiquitously in human hair, breast milk, and blood and in environmental populations, such as water basins, surface soils, and plants cultivated in contaminated sites (4–10). Exposure to PFAS has been linked to several adverse human health effects, including immunotoxicity, liver damage, and thyroid disfunction and negative ecological impacts of endocrine disruption in aquatic species (11–14).

To date, the U. S. Environmental Protection Agency (USEPA) and the international Organization of Economic Co-operation and Development (OECD) have categorized thousands of individual compounds based on a threshold of fluorine percentage and molecular structure or the presence of perfluoroalkyl moieties (15, 16). The structures of these compounds are distinct with linear and branched carbon chains of varying length and fluorination and functional head groups containing sulfonates, sulfonamides, carboxylates, and other moieties that are believed to govern compound stability, transport, and environmental fate (17). In addition to the variety of PFAS across industrial products, environmental samples from a single contaminated source will often contain mixtures of intact PFAS, partial degradation products, and precursor chemicals (18, 19). Sample complexity and batch variation make measurement and quantitation of PFAS-containing samples with traditional analytical techniques like liquid chromatography – mass spectrometry (LC-MS) (20–22) challenging and costly. Moreover, water remediation strategies including ion exchange resins, high-pressure membrane systems, and granular activated carbon (23) and soil remediation strategies including thermal treatment and soil washing (24) are also complicated by the unique chemical properties that originally made PFAS efficient in industrial and commercial products. To rapidly respond to this growing problem, there is a need for alternative approaches to real-time environmental monitoring and mitigation efforts, as well as comprehensive methods to assess environmental impact (25–27). Increasingly, researchers are turning to biology as an alternative approach for PFAS detection (28, 29) and mitigation (25, 30). Broad measurements of living systems could not only aid in rapid, on-site detection of new contamination sources and identification and classification of new PFAS byproducts but also expand our general understanding of the biological consequences of PFAS contamination (25, 26). Bacteria are a promising measurement platform as they play critical roles in most ecosystems, are highly concentrated in soil environments and aquifers where PFAS contamination persists (31, 32), and rapidly sense and respond to environmental fluctuations (33, 34). Transcriptomic analysis by RNA sequencing (RNA-seq) of bacterial interactions with PFAS could serve as an alternative measurement strategy to explore genome-wide molecular mechanisms underlying gene expression level changes induced by PFAS. RNA-seq is a widely applicable technique that can investigate the response of a wide variety of organisms to various types and mixtures of PFAS, potentially enabling the identification of specific sensing gene targets and PFAS degradation pathways and provide insight into how these non-natural chemicals interact with living systems. Although, omics-based analyses have been used to examine the biological response of several organisms, including primary human liver cell spheroids, zebrafish, and *Escherichia coli*, to various PFAS (35–38), studies exploring the global transcriptional response of bacteria across key growth phases are limited.

In this study, we employed time course RNA-seq at key phases of bacterial growth to investigate the global transcriptional response of a model Gram-negative bacterium, *E. coli* MG1655, to PFAS (Figure 1). *E. coli* is an optimal organism to begin understanding the microbial transcriptional responses to PFAS, as it is present in different environments and its transcriptional responses to numerous stressors and growth conditions have been extensively studied with RNA-seq (39–42). *E. coli* was grown in the presence of perfluoroalkyl carboxylic acids (PFCA) or equivalent non-fluorinated carboxylic acids (NFCA) at a concentration of 100 μmol/L for 48 hr, and samples were sequenced at mid-exponential and stationary phase. Differential expression analysis revealed significant changes in mid-exponential and stationary phase gene expression induced by two different PFCAs, including changes that differed from the NFCAs. Our results indicate *E. coli* employed different mechanisms at different growth phases further signifying temporal gene expression in response to PFCAs. Specifically, we identified significant changes in the expression of the formate regulon and sulfate assimilation at mid-exponential phase, ferrous iron transport, amino acid metabolism, small regulatory RNAs during early stationary phase, and central metabolism, ribosome biosynthesis, molecular chaperones, sigma factors, and motility process of flagellar assembly and chemotaxis during late stationary phase. The transcription-level responses of *E. coli* to PFCAs presented in this study expand our knowledge of the molecular-level response to PFAS and could inform the development of future environmental monitoring and mitigation strategies.

**Figure 1.**
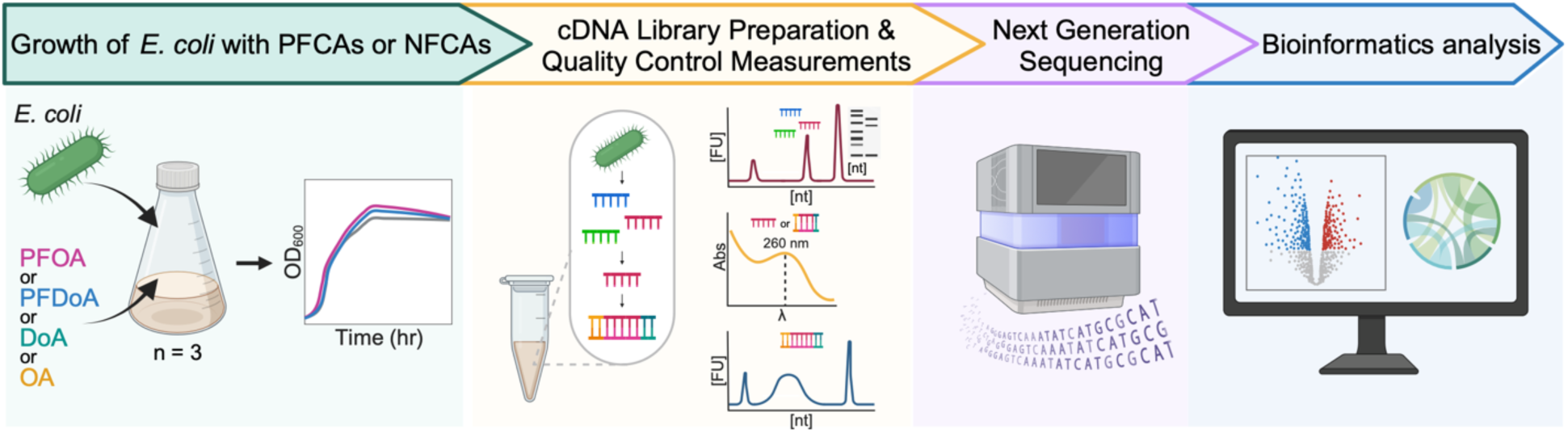
Experimental and computational workflow. This study characterizes the PFCA-induced transcriptional response of *E. coli* during mid-exponential phase and stationary phase growth. The schematic diagram of experimental design and data processing provides an overview of the growth of *E. coli* with PFCAs or NFCAs, microbial growth measurements, and cDNA library preparation and quality control measurements before next generation sequencing, bioinformatic data processing and differential expression analysis. Additional culturing and sampling information can be found in SI Figure 1.

## RESULTS AND DISCUSSION

### Culturing *E. coli* with 100 μmol/L of PFCA does not inhibit growth

To study the transcriptional responses of *E. coli* to PFAS, we first needed to understand the growth characteristics of *E. coli* in the presence of PFAS. From a sub-class of PFAS, two PFCAs, perfluorooctanoic acid (PFOA) and perfluorododecanoic acid (PFDoA), were selected as representative PFCAs commonly found in environmental contamination (12, 43) and to provide a comparison of chain length of PFCAs between 8 and 12 carbons, respectively (Figure 2 A). We aimed to select a concentration high enough to induce a measurable change in gene expression but low enough to avoid toxicity to the cells. Based on existing literature (37, 44) and the observation that *E. coli* cell division and growth can stall with fluoride concentrations between 10 μmol/L and 100 μmol/L (45–47), we initially selected these concentrations to explore PFOA and PFDoA toxicity. *E. coli* was grown in M9 minimal medium supplemented with glucose and 10 μmol/L or 100 μmol/L of either PFOA or PFDoA for a period of 48 hours at 37 °C (SI Figure 1, Materials and Methods). The PFCAs were dissolved in dimethyl sulfoxide (DMSO), so to generate untreated control samples for differential expression analysis, cells were cultivated in the same liquid medium with an equal concentration of DMSO not containing PFCAs. NFCAs, octanoic acid (OA) and dodecanoic acid (DoA) (Figure 2 A), were also included to compare the response of *E. coli* to fluorinated and non-fluorinated carboxylic acids. All sample groups were prepared in biological triplicate (SI Figure 1). We monitored growth of *E. coli* in the presence and absence of PFCAs or NFCAs using optical density measurements at 600 nm (OD_600_) at select time points (Figure 2 B; SI Figure 2) across 48 hours. *E. coli* exhibited similar growth profiles for both 100 µmol/L (Figure 2 B) and 10 µmol/L (SI Figure 2) of PFOA, PFDoA, OA, and DoA, and a two-way mixed ANOVA confirmed no statistically significant difference in growth to the untreated control (Table SI 1). As both concentrations tested yielded similar growth profiles, we elected to collect samples for sequencing and downstream analysis from cells grown with 100 µmol/L of each compound.

**Figure 2.**
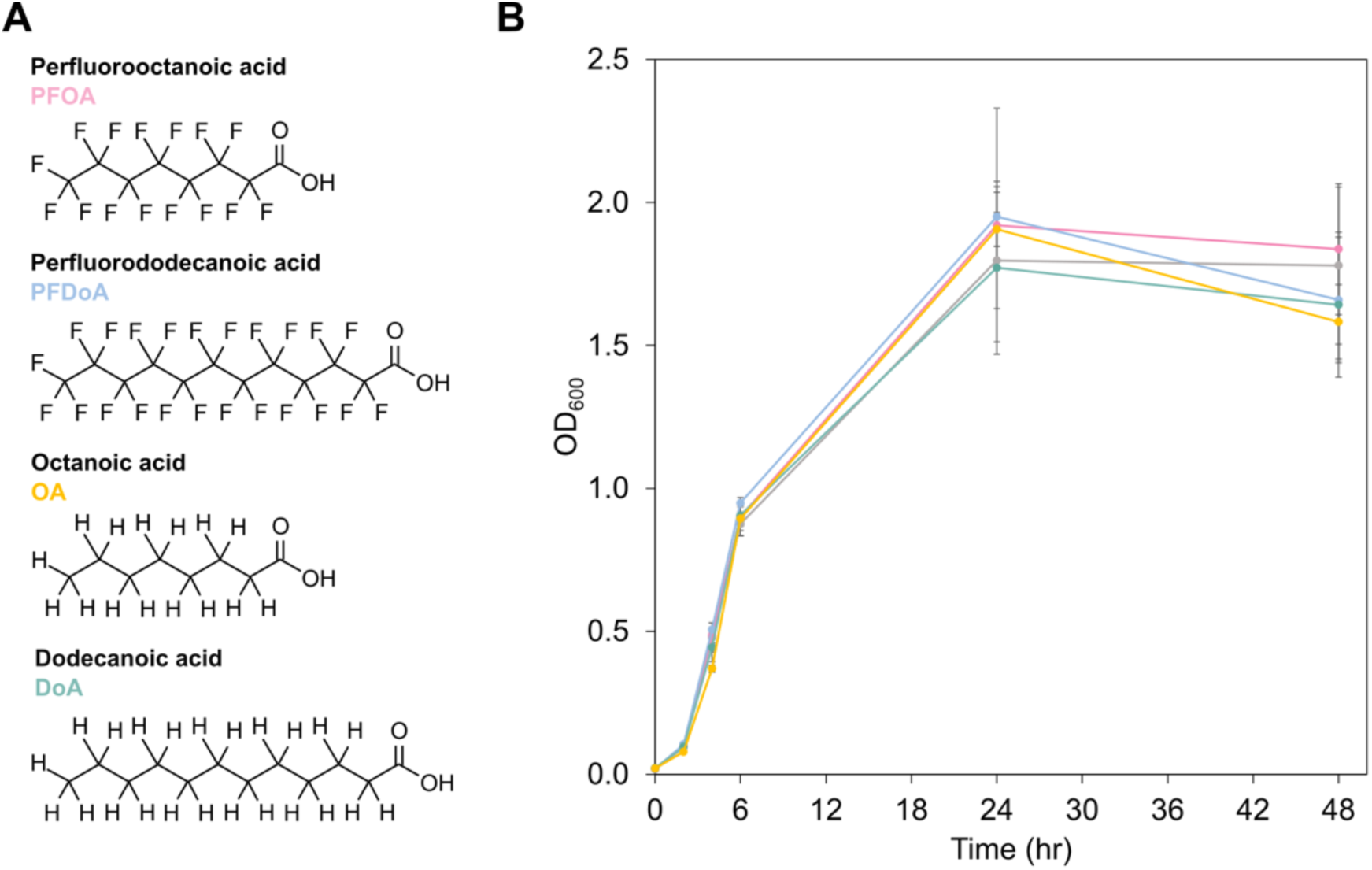
Growth of *E. coli* in the presence of PFCAs or NFCAs. (A) Chemical structures of PFOA, PFDoA, OA, and DoA. (B) Growth of *E. coli* in M9 at 37 °C in the presence of 100 μmol/L of PFOA (pink), PFDoA, (blue), OA (yellow), or DoA (green) displayed as optical density measured at 600 nm (OD_600_) over time (hr). Untreated cells containing an equivalent final concentration of DMSO are displayed in gray. Error bars represent one standard deviation of the mean value of biological triplicates (Table SI 1).

### Global transcriptional response of *E. coli* to PFCAs and NFCAs across the time course

Next, we collected samples for total RNA extraction and next generation sequencing (NGS) at 6 hr, 24 hr, and 48 hr. Sampling times were selected to represent mid-exponential phase, the transition from exponential to stationary phase, and the end of stationary phase of bacterial growth. Gene expression naturally shifts between these growth phases (48); therefore, we hypothesize PFCAs will further exaggerate the natural changes in expression. We extracted total RNA from three biologically distinct cultures of each sample type for commercial cDNA library preparation and sequencing (Materials and Methods). NGS generated a total of over 350 million reads with a mean quality score of 35.88 (Table SI 2).

We performed principal component analysis (PCA) of normalized data through DESeq2 methodology (49) to visualize variation in gene expression between the 45 samples in this study (SI Figure 3). All transcriptomes first cluster with their respective growth phase as seen with PC1, implying an inherent temporal transcriptomic response. Transcriptomes of 24 hr and 48 hr samples had less variation and clustered closer to one another than to the 6 hr transcriptomes, suggesting gene expression during the two points within stationary phase was more similar than mid-exponential phase at the time of sampling. These results demonstrate the global shift in gene expression is governed by a shift from active growth to cell survival and response to stress regardless of sample type. Additionally, PFOA samples clustered more closely to PFDoA samples than the untreated control samples throughout the time course (SI Figure 3), indicating additional difference in gene expression was likely induced by PFCAs. At 6 hr and 24 hr, NFCA transcriptomes clustered more closely to untreated control transcriptomes than PFCA transcriptomes, suggesting gene expression of NFCAs and untreated samples were more similar compared to PFCA samples. PCA indicated a biological replicate of DoA at 48 hr had a large variation in gene expression from the triplicate group (outlier); as a result, DoA triplicate samples at 48 hr were removed from further analysis.

We employed differential gene expression analysis to determine if PFCAs induced a measurable change in *E. coli* gene expression over the time course. In this study, we consider genes to be differentially expressed if the adjusted p-value (padj) is less than 0.05. Additionally, we mark moderate and significant differential expression with absolute values of the log_2_ fold change greater than or equal to 1.2 and 1.5, respectively. Untreated samples serve as the reference level for gene expression pairwise comparisons. The global transcriptional response of *E. coli* to PFOA and PFDoA demonstrates a measurable change in gene expression was induced by both PFCAs throughout the time course (Figure 3 A, B). A similar number of genes were differentially expressed by PFOA and PFDoA at 6 hr and 24 hr (Figure 4 A). However, this trend diverges at 48 hr, as PFOA induced the most differentially expressed genes and PFDoA induced the least. The considerable change in gene expression induced at 48 hr by PFOA could be due to many factors, which we will explore in more detail in later sections. NFCAs also induced a measurable change in gene expression when compared to the untreated control across the time course (Figure 4 B; SI Figure 4 A, B). *E. coli* grown with OA exhibited a similar pattern of expression to PFOA at 6 hr and 24 hr (SI Figure 5 A). However, PFOA induced differential expression of more than 700 genes at 48 hr, in contrast, OA induced less than 40 genes (SI Figure 5 B), indicating differed response to fluorinated and non-fluorinated versions of the 8-chain compound. DoA had an opposite result from PFDoA, inducing differential expression in over 350 genes compared to over 50 genes at 6 hr (Figure 4 A, B; SI Figure 5 C), further suggesting differed responses to fluorinated and non-fluorinated versions of the 12-chain compound.

**Figure 3.**
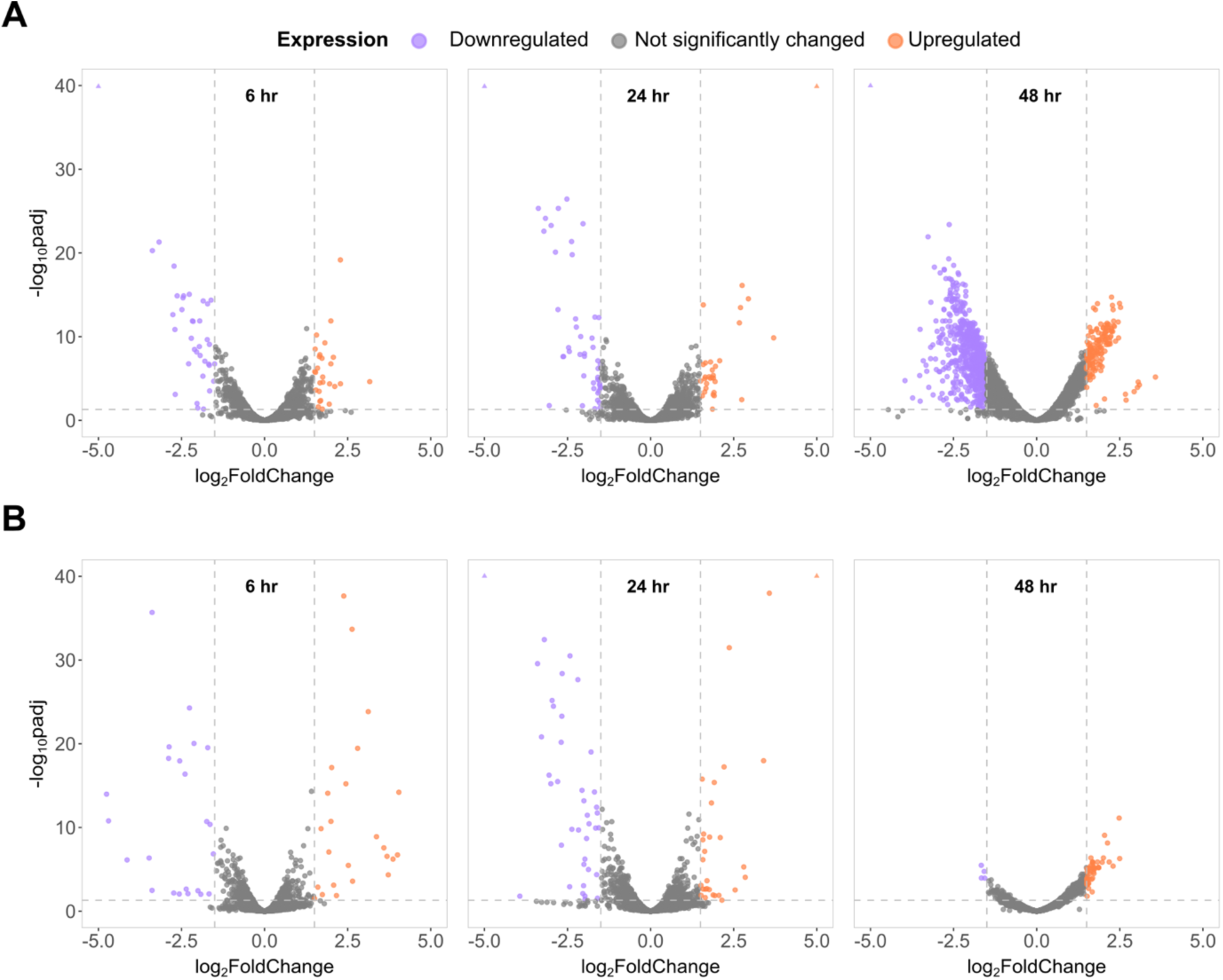
Transcriptomic profiles of mid-exponential and stationary phase *E. coli* grown with PFCAs. Volcano plots show the global transcriptional response of *E. coli* to (A) PFOA and (B) PFDoA in comparison to an untreated control group at 6 hr (left), 24 hr (middle), and 48 hr (right). Dashed lines represent a significance cutoff for differential expression of an adjusted p-value (padj) of less than 0.05 and absolute value of the log_2_ fold change greater than or equal to 1.5. Upregulated and downregulated genes are denoted by purple and orange, respectively, and genes not significantly changed by PFCA treatment are gray. Differentially expressed genes with log_2_ fold change and or padj values outside of the plot limits are represented by orange or purple triangles in the upper corners of each plot, if applicable.

**Figure 4.**
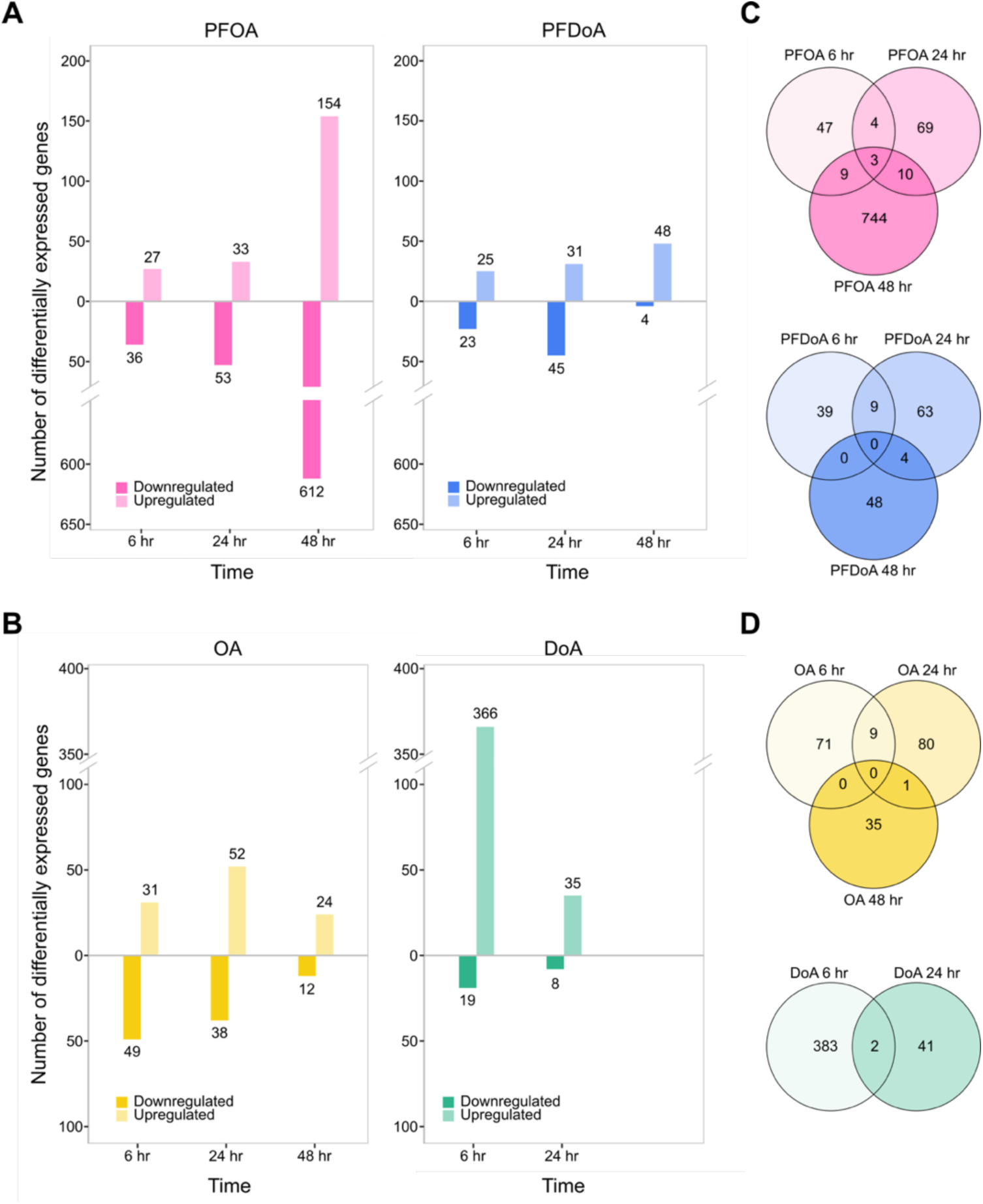
Comparison of differential gene expression of *E. coli* induced by PFCAs or NFCAs across the time course. The number of differentially expressed genes induced by (A) PFOA (pink, left), PFDoA (blue, right), (B) OA (yellow, left), and DoA (green, right) over the time course are shown by the height of each bar and label above or below the bar. Genes are considered differentially expressed if the absolute value of the log_2_ fold change greater than or equal to 1.5 and the adjusted p-value less than 0.05. Upregulated genes are represented by the upper portion of the plot with lighter shaded bars as indicated by the legend. Downregulated genes are represented by the lower portion of the plot with darker shaded bars. Venn diagrams display the number of differentially expressed genes induced by (C) PFOA (top), PFDoA (bottom), (D) OA (top), and DoA (bottom) over the time course. Unions display genes differentially expressed by one PFCA or NFCA at more than one time point. Genes in the outer circles are only differentially expressed at a single time point by one PFCA or NFCA.

We observed a temporal pattern to PFCA- and NFCA-induced differential expression, as the majority of genes were significantly changed at only one time point (Figure 4 C, D). Less than 30 and 15 genes were differentially expressed at multiple time points by PFOA and PFDoA, respectively (Table SI 3; Table SI 4). Only three genes*, gatD, glpA*, and *yeaR,* which are related to galactose metabolism and transport (50) and glycerol metabolism (51), were differentially expressed by PFOA at all time points in the study (Figure 4 C top; Table SI 3). Genes encoding formate hydrogenlyase subunits, iron transporters, and small regulatory RNAs were differentially expressed by PFOA at two of the three time points and will be discussed in later sections. Unlike PFOA, no genes were differentially expressed in *E. coli* by PFDoA at all time points (Figure 4 C bottom). Upon further investigation, nine genes, including the *trp* operon, downregulated by PFDoA at 6 hr and 24 hr, were also downregulated by PFOA and OA at 24 hr and DoA at 6 hr (SI Figure 5 D, E; Table SI 5), and will be discussed in a later section. Also discussed later, three of the four genes upregulated by PFDoA at 6 hr and 24 hr were also downregulated by PFOA at 24 hr and 48 hr and OA at 6 hr (Table SI 5). These genes compose the ferrous iron transport operon, *feo*ABC. Similar to PFOA, we observed a temporal pattern of OA-induced differential expression over the time course with minimal common expression between time points (Figure 4 B, D top; Table SI 6). Throughout the time course, PFOA and OA induced 13, 32, and 35 of the same genes at 6 hr, 24 hr, and 48 hr, respectively (SI Figure 5 A, B; Table SI 7). These results suggest responses were triggered by the octanoic acid moiety regardless of fluorination. Additionally, DoA commonly induced differential expression in two genes, *prpC* and *ydeO*, involved in propionate catabolism (52) and regulation of the *gad* system (53), at 6 hr and 24 hr (Figure 4 D bottom; Table SI 8), similar to minimal shared expression of 9 genes by PFDoA at 6 hr and 24 hr (Figure 4 C bottom; SI Figure 5 C; Table SI 4). Parallel to common expression between PFOA and OA, PFDoA and DoA induced differential expression in the same 19 genes at 6 hr and 13 other genes at 24 hr (SI Figure 5 C; Table SI 7). These results indicate differential expression induced by PFCAs or NFCAs is predominantly specific to a particular phase of growth.

### Transcriptional response of mid-exponential *E. coli* to PFCAs

Bacterial gene expression shifts significantly between exponential phase and stationary phase, reflecting the transition from active growth and energy production towards stress-mitigation and survival in resource-limited environments (48, 54). We hypothesize PFCAs would alter gene expression differently during exponential phase and stationary phase. To better understand the response of *E. coli* to PFCAs at each growth phase within the study’s time course, we sought to identify genes that were differentially expressed by PFOA and PFDoA (common responses) and the genes only changed by one PFCA (specific responses), starting at mid-exponential phase (6 hr). Highlights of our results of the response of *E. coli* to PFCAs and NFCAs across the time course, discussed below, are illustrated in Figure 5.

**Figure 5.**
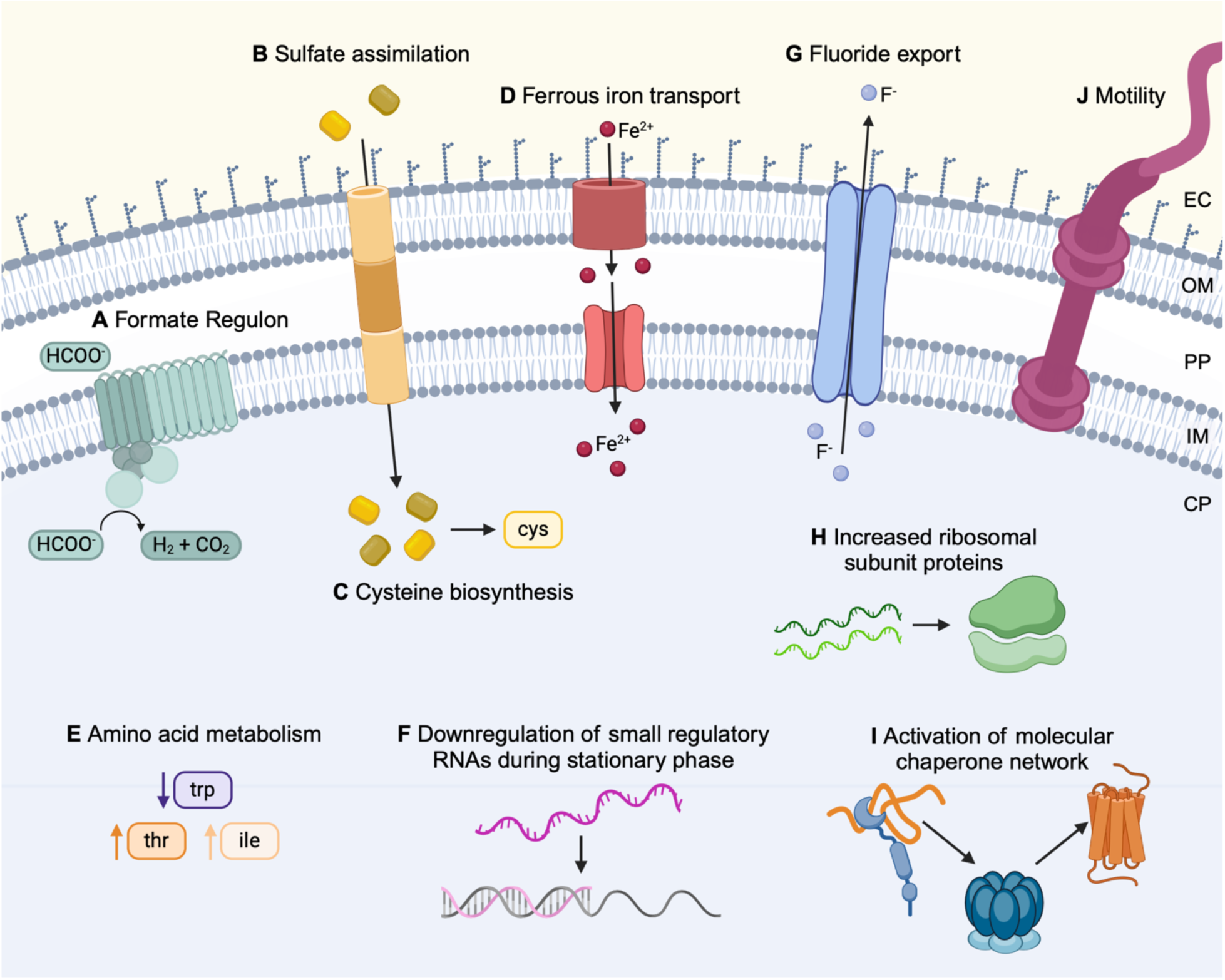
Schematic of key biological processes with altered gene expression in the presence of PFCAs across the time course. These processes are highlighted in the results and discussion. The formate regulon (A), sulfate assimilation (B), and cysteine biosynthesis (C) are discussed in the mid-exponential phase section. Ferrous iron transport (D), amino acid metabolism (E), and small regulatory RNAs (F) are discussed in the early stationary phase section. Fluoride export (G), ribosomal subunit proteins (H), the molecular chaperone network (I), and motility processes (J) are discussed in the late stationary phase section. The cytoplasm (CP), inner membrane (IM), periplasm (PP), outer membrane (OM), and extracellular environment (EC) are labeled on the right side of the schematic. The CP, PP, and EC have blue, white, and yellow backgrounds, respectively.

PFOA and PFDoA each induced differential expression of approximately 50 genes at 6 hr (Figure 4 A; Table SI 3; Table SI 4). This is less than 2% of genes in the transcriptome, indicating growth in the presence of PFCAs does not seem to have a major impact on gene expression of mid-exponential *E. coli*. Only seven genes were differentially expressed by both PFCAs at 6 hr (SI Figure 5 D). These genes, *nadA*, *msyB*, *hiuH*, *galS*, *adiY, yceK,* and *ymfQ,* encode a quinolinate synthase, acidic protein, hydroxyisourate hydrolase, DNA-binding transcriptional dual regulator and activator, and two uncharacterized proteins, respectively (Table SI 5). With roles in pyrimidine and purine metabolism, quinolinate synthase and hydroxyisourate hydrolase were also downregulated by PFOA and PFDoA at 24 hr, indicating a sustained effect on gene expression. As an (4Fe-4S) cluster is required for activity of quinolinate synthase in NAD synthesis (55), downregulation of *nadA* could suggest a requirement for iron and sulfur that correlates with upregulation of sulfate assimilation and ferrous iron transport genes discussed in later sections. The DNA-binding transcriptional dual regulator GalS was upregulated by PFOA at 6 hr, but downregulated by PFDoA, suggesting *E. coli* exhibited a different response to the 8-chain PFCA compared to the 12-chain. Several of these genes were also downregulated in response to the NFCAs (Table SI 5; Table SI 7). For example, significant downregulation of *msyB* by both PFCAs and NFCAs implies some changes in gene expression were not specific to fluorinated compounds, but likely rather to carboxylic acids. The minimal common changes in gene expression at 6 hr implies *E. coli* utilized different mechanisms at mid-exponential phase to respond to PFOA and PFDoA.

During mid-exponential phase, *E. coli* differentially expressed several genes in response to PFOA but not PFDoA, including the formate regulon, a putative transporter YhjX, L-aspartate oxidase, L-alanine exporter, and small RNA RyjA (Table SI 3). Of particular interest, genes encoding components of the formate regulon, including the formate hydrogenlyase (FHL) complex, formate dehydrogenase H, and the hydrogenase pleiotropic *hyp* operon (56), were significantly downregulated (Figure 5 A; Figure 6 A; Table SI 9). The FHL complex is essential to fermentative H_2_ production in *E. coli* during mixed-acid fermentation. At low extracellular pH and high levels of re-imported formate during stationary phase, the FHL complex is activated to disproportionate accumulated formate to hydrogen and carbon dioxide (57, 58). We found the formate channel, encoded by *focA,* that transports formate from the cytoplasm during exponential phase under physiological pH, was not differentially expressed at 6 hr by either PFCA or NFCA, indicating these compounds did not induce more or less formate transport than the untreated control. However, formate dehydrogenase H, encoded by *fdhF,* and the *hyc* operon, composed of genes encoding the cytoplasmic domain subunits, *hycBEFG*, and membrane domains, *hycCD*, of the FHL complex, plus the transcriptional regulator, *hycA*, chaperone, *hycH*, and protease, *hycI*, were downregulated by PFOA at 6 hr (Figure 6 A; Table SI 9). Additionally, *hycBCDE* were also significantly downregulated by PFOA at 48 hr, indicating the change in expression of these four genes was not unique to mid-exponential phase. Four genes of the *hyp* operon, *hypABCD*, encoding products involved in nickle:iron hydrogenase maturation (56, 59), and a gene encoding a putative transporter predicted to function as an oxalate:formate antiporter, *yhjX* (60, 61), were also downregulated at 6 hr by PFOA but not differentially expressed by PFDoA or either NFCA. Downregulation of formate regulon genes, unfavorable reaction conditions in an aerobic environment, and the unlikely occurrence of common inducing factors like acidic pH, the absence of nitrates, and unlikely PFOA-induced formate accumulation during mid-exponential phase, suggests forward conversion of formate was possibly prevented (57). These results further signify a specific response to PFOA that differs from routine transport and conversion of formate under physiological conditions. Lastly, formate is a degradation product of PFOA, and additional measurements would be needed to determine if formate ions from PFOA were present in the cultures.

**Figure 6.**
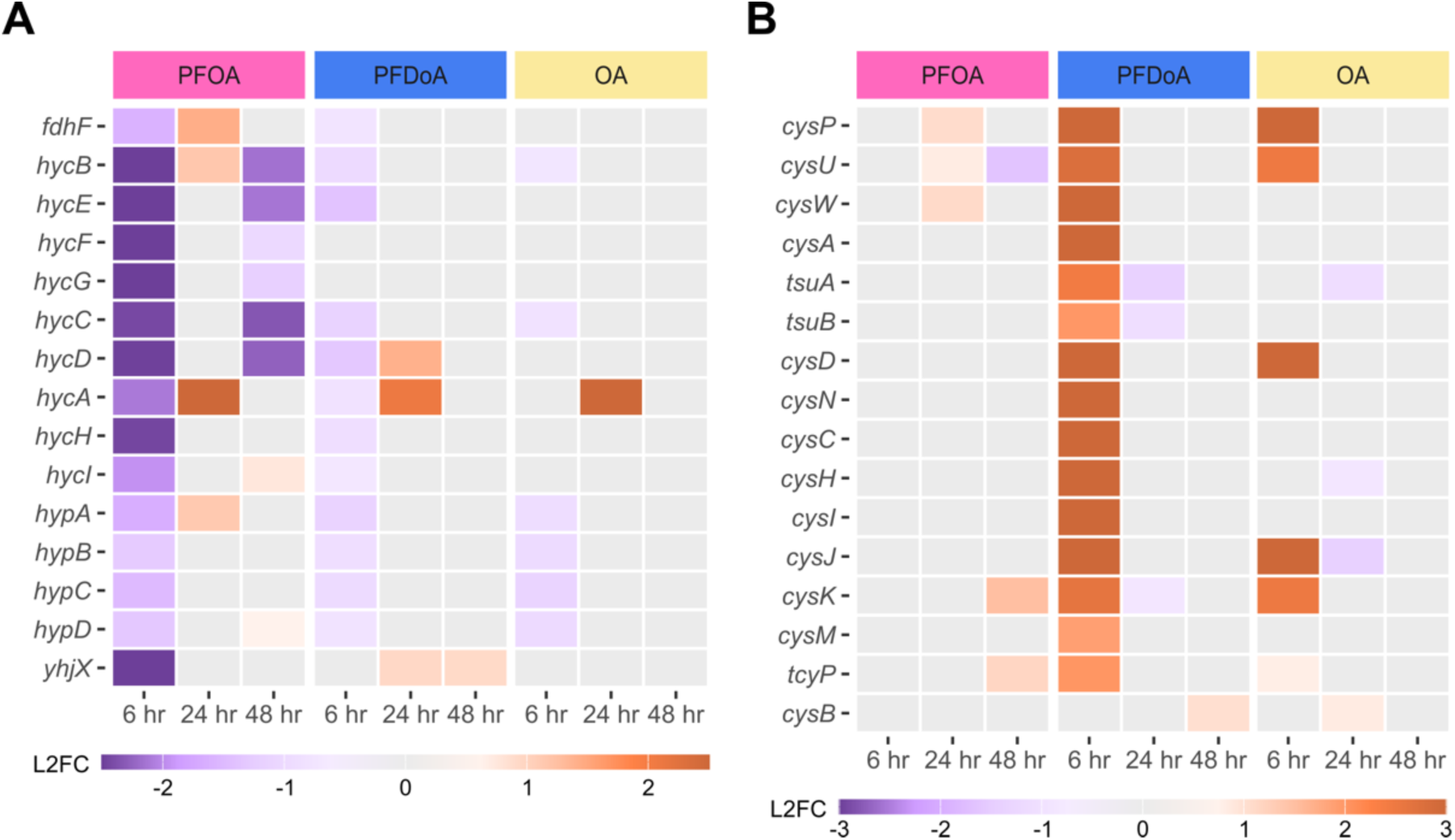
Select genes with differential expression at 6 hr and their temporal change. Heatmaps display differential expression of select genes (padj < 0.05) involved in (A) the formate regulon and (B) sulfate assimilation and cysteine biosynthesis. Expression is represented by log_2_ fold change (L2FC) within a gene compared to untreated control in a purple to orange scale displayed by legend color bar below each heatmap. Note heatmap color bars are different scales in (A) and (B) to best highlight differences. PFCA or NFCA treatment is represented by color coded bars at the top of the heatmaps: PFOA (pink), PFDoA (blue), and OA (yellow). Genes with no statistically significant change in expression are represented by gray boxes.

During mid-exponential phase, PFDoA induced overexpression of genes involved in sulfate and thiosulfate assimilation and L-cysteine biosynthesis (Figure 5 B, C; Figure 6 B; Table SI 4; Table SI 10). Sulfate assimilation is crucial for the growth and survival of *E. coli*, as sulfur is needed to synthesize cysteine and many cofactors and is important for cellular redox balance (62). Sulfate and thiosulfate transporters are often overexpressed in *E. coli* when intracellular sulfur is limited, and uptake of extracellular sulfate is needed (63). Newly assimilated sulfur-containing intermediates are then converted to cysteine (64). The sulfate/thiosulfate ATP-binding cassette (ABC) transporter complex, *cysPUWA,* was significantly upregulated with log_2_ fold changes greater than 2.75 by PFDoA at 6 hr (Figure 6 B; Table SI 10). This was not observed with PFOA or DoA. Genes encoding the transporter’s periplasmic protein, *cysP*, and inner membrane subunit, *cysU*, were also significantly upregulated by OA at 6 hr, indicating possible uptake of sulfate was induced by OA as well (SI Figure 5 D). In a second path for sulfur assimilation, genes, *tsuAB*, in a recently characterized thiosulfate ion uptake mechanism (65), were also upregulated by PFDoA at 6 hr (Figure 6 B), but were not significantly changed by DoA. These results suggest PFDoA induced a specific response to utilize this mechanism for thiosulfate ion uptake, and the response was likely triggered by the 12-chain PFCA. Following sulfate and thiosulfate uptake, the genes encoding enzymes that catalyze the activation and subsequent reduction of sulfate to intermediates, sulfite and sulfide, were upregulated by PFDoA. The sulfate activation operon, *cysDNC*, responsible for chemical activation of sulfate to adenosine 5’-phosphosulfate (APS) by ATP sulfurylase and APS kinase and the gene cluster, *cysHIJ*, encoding APS reductase and sulfite reductase subunits, responsible for the reduction of APS to sulfite and sulfite to sulfide (66, 67), respectively, were overexpressed by PFDoA (Figure 6 B).

In addition to upregulation of sulfur-containing compound transporters and redox enzymes, unsurprisingly, cysteine synthases, *cysKM* (64), and an inner membrane L-cystine (dimeric cysteine) transporter, *tcyP*, were significantly upregulated by PFDoA at 6 hr (Figure 5 B; Figure 6 B). Furthermore, when grown on minimal medium, as done in this study, TcyP functions as the main importer of L-cystine (68) and is activated when intracellular cysteine is low (64). These changes in gene expression could indicate an increase requirement for cysteine through sulfate assimilation or direct import in response to PFDoA. Lastly, the gene encoding the regulatory protein CysB, which drives the sulfur limitation response in *E. coli,* was not differentially expressed by any PFCA or NFCA treatment in this study (66, 69). It is possible the gene was overexpressed at an early point in exponential phase as the genes it regulates were overexpressed by PFDoA at the mid-exponential mark or has consistent expression. Overexpression of sulfate and thiosulfate assimilation components and cysteine biosynthesis enzymes by PFDoA, indicates PFDoA could be negatively impacting redox balancing and in turn, stimulating an increased requirement for sulfur-containing compounds like cysteine during mid-exponential phase.

### Transcriptional response of early stationary phase *E. coli* to PFCAs

During stationary phase, bacteria modulate gene expression by simultaneously activating essential genes for survival and stress mitigation, while preventing transcription of unnecessary genes in a nutrient-deficient environment. Additional stress induced by the prolonged presence of PFCAs may further modify gene expression during stationary phase when compared to mid-exponential phase. Unlike mid-exponential phase, the number of genes differentially expressed specifically by PFOA or PFDoA during stationary phase at 24 hr was minimal (SI Figure 5 D, E). PFOA induced specific changes in gene expression of genes encoding cytochrome bo3 subunits, biotin biosynthesis enzymes, and transporters for iron and tryptophan (Table SI 3), whereas PFDoA induced changes in gene expression of inner membrane proteins and the small regulatory RNA GadY (Table SI 4). In contrast to minimal common responses observed at 6 hr, PFOA and PFDoA induced differential expression of the same 49 genes at 24 hr (SI Figure 5 D, E, F; Table SI 5). Products of these genes have roles in ferrous iron transport (Figure 7 A), amino acid metabolism (Figure 7 B), and small RNA regulation (Figure 7 C). Common changes in gene expression suggest these responses are not specific to a particular length of PFCAs, but to the PFCAs explored within the scope of this study. Expression of 24 of the 49 genes were also significantly changed by OA at 24 hr and only 10 were changed by DoA (SI Figure 5 E; Table SI 5; Table SI 7), indicating those genes likely have a larger role regarding carboxylic acids in general, rather than fluorination.

**Figure 7.**
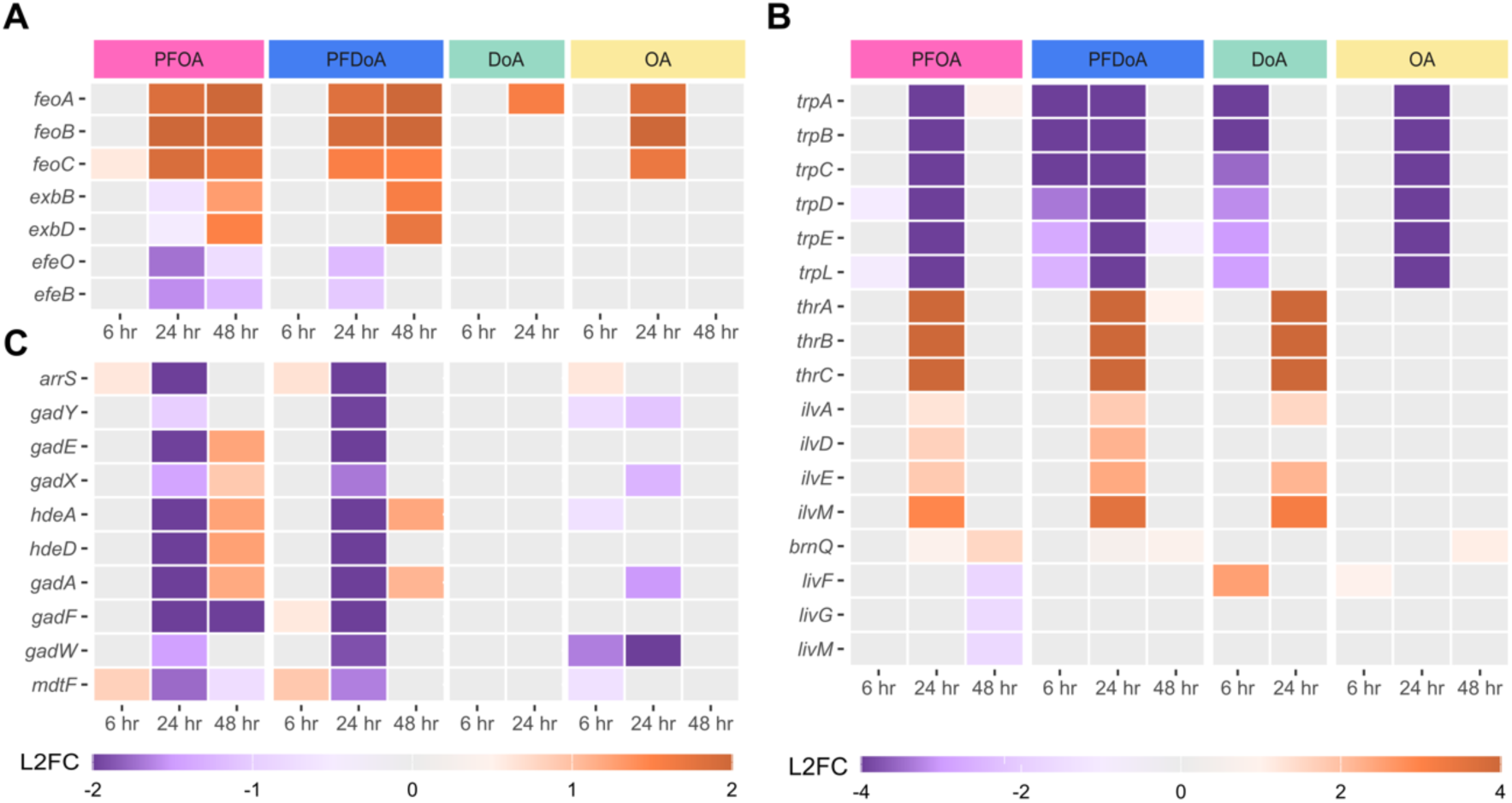
Select genes with differential expression at 24 hr. Heatmaps display differential expression of select genes (padj < 0.05) involved in (A) ferrous iron transport, (B) amino acid metabolism, and (C) small regulatory RNAs across the time course. Expression is represented by log_2_ fold change (L2FC) within a gene compared to untreated control in a purple to orange scale displayed by legend color bar below each heatmap. Note panels A and C share the same L2FC scale, which differs from panel B. PFCA or NFCA treatment is represented by color coded bars at the top of the heatmaps: PFOA (pink), PFDoA (blue), DoA (green), and OA (yellow). Genes with no significant change in expression are represented by gray boxes.

As iron is an essential nutrient and cofactor in various enzymes, several iron acquisition systems and regulators have evolved to maintain intracellular iron homeostasis (70, 71). Our analysis revealed genes encoding components of iron uptake transport mechanisms were differentially expressed in response to PFCAs (Figure 5 D; Table SI 5; Table SI 11). One operon *feoABC*, which encodes ferrous iron (Fe^2+^) transporters (72), was significantly upregulated at 24 hr and 48 hr by both PFOA and PFDoA (Figure 7 A; SI Figure 5 F). The operon was also upregulated at 24 hr by OA (Table SI 11), implying the elevated response could be stimulated by the carboxylic acid moiety. Upregulation of *feo,* in combination with no significant change in the expression of upstream Fur and Fnr transcripts, the ferric iron (Fe^3+^) transcriptional regulator and activator, (73, 74), suggests PFCAs induce a requirement for intracellular ferrous iron during stationary phase. We hypothesize this is due to preference for the soluble ferrous form in comparison to insoluble ferric form at the pH of the culture. Moreover, the level of free iron could increase via *feo* transport because a putative Fur-regulated inhibitor, *rhyB,* (75, 76) was downregulated by PFOA at 48 hr. Additionally, fluoride exposure can deplete active forms of metal ions, including iron, through formation of stable metal-fluoride complexes, therefore, triggering cells to increase free metal ion concentration (77, 78) to maintain homeostasis. Siderophores, secreted ferric iron scavengers (79, 80), were not employed because the enterobactin gene cluster and associated transporters were moderately downregulated at 48 hr by PFOA (Table SI 11). However, genes, *exbB* and *exbD*, in the associated TonB energy-transducing complex (81) were upregulated by PFOA and PFDoA at 48 hr (Figure 7 A). The *efeUOB* operon encoding a ferrous iron transporter complex is not fully functional in *E. coli* MG1655 due to a frameshift mutation in *efeU* (82), but the genes encoding periplasmic proteins*, efeO* and *efeB*, were also downregulated by both PFCAs at 24 hr. Additionally, PFOA-induced downregulation of genes encoding various cytochromes (Table SI 11) during stationary phase could suggest limited available iron for incorporation into heme groups (83–85). These results further signify a specific preference and need for ferrous iron uptake by the *feo* operon during stationary phase and likely limited availability for incorporation in key components like cytochromes (84).

During stationary phase, gene expression is altered to meet metabolic requirements including the adjustment of amino acid metabolism (86, 87). Genes encoding the *trp* operon, the *thr* operon, isoleucine synthesis components (*ilvADE* and *ilvM*), and transporters of branched-chain amino acids (*brnQ* and *livFGM*) were differentially expressed by PFOA and PFDoA during stationary phase (Figure 5 E; Figure 7 B; Table SI 11). Notably, differences in gene expression between PFCA samples and the untreated controls of the *trp* and *thr* operons were some of the largest in this study, with log_2_ fold changes less than -5.3 and greater than 4.6, respectively. Downregulation of the *trp* operon would suggest an effort to route metabolic flux towards optimal parts of central metabolism or amino acid biosynthesis pathways (SI Figure 6) (88), in order to adapt to high levels of intracellular tryptophan or related PFCA-induced stress. Alternatively, upstream precursors to the *trp* operon, like chorismate (89), may not have been available, and as a result, the operon was underexpressed. The *trp* operon was also downregulated by DoA at 6 hr and OA at 24 hr (Figure 7 B; Table SI 11), indicating the change in gene expression was possibly prompted by the octanoic acid moiety regardless of the presence or absence of fluorination. Moreover, downregulation of tryptophan symporters, *mtr* and *tnaB*, at 24 hr and no change in expression of the general aromatic acid transporter, *aroP*, imply extracellular tryptophan was likely not imported into cell (Table SI 11). Upregulation of threonine and isoleucine biosynthesis genes could be a direct result of carbon flux redirection from upper metabolism, and the synthesized amino acids (90, 91) could be used in processes to mitigate stress and sustain growth during nutrient-limited stationary phase.

Small regulatory RNAs (sRNAs) help modulate gene expression during stationary phase and under environmental stress by stabilizing specific mRNAs and stimulating translation (92). At 24 hr, 10 sRNAs, their corresponding mRNAs targets, and flanking genes were downregulated by PFOA and PFDoA (Figure 5 F; Figure 7 C; Table SI 11). A handful of these genes remained downregulated by PFOA at 48 hr. In this study, this response was fairly specific to PFCAs, as only four of these genes were differentially expressed by NFCAs. Also, the genes encoding RNA-binding proteins Hfq and ProQ were minimally overexpressed by PFOA at 48 hr with log_2_ fold changes of approximately one. These proteins work in tandem with sRNAs to bind target mRNAs and provide intracellular stability (93). Downregulation of antisense sRNAs genes, *arrS* and *gadY*, the corresponding mRNAs that they target, *gadE* and *gadX*, and flanking genes, *hdeAD, gadAFW, and mdtF,* during stationary phase by PFOA and PFDoA (Figure 7 C), suggests possible energy conservation and resource reallocation to critical survival pathways (92, 94) in response to PFCAs. Lastly, *arrS* and *gadY* products have essential roles in acid resistance regulation (94), and downregulation of these sRNAs at 24 hr suggests PFCAs either do not significantly induce acid stress compared to the untreated control, or the activation of pH homeostasis mechanisms is too costly due to nutrient limitation during stationary phase.

### Transcriptional response of late stationary phase *E. coli* to PFCAs

To compare the transcriptional response of *E. coli* to PFCAs at different points within stationary phase, we wanted to understand how differential expression patterns induced at 48 hr differed from patterns at 24 hr. At 48 hr, PFOA and PFDoA upregulated 28 common genes that encode the 50S ribosomal subunits, the DNA-binding transcriptional repressor LsrR, and Ton complex subunit ExbD (Table SI 5; SI Figure 5 E, F). Seventeen of these genes were also moderately upregulated by OA (Table SI 7), with a log_2_ fold change of at least 1.4, indicating differential expression may not be specific to PFCAs. We observed specific changes in gene expression of cold shock proteins and putative phosphotransferase system enzymes by PFDoA at 48 hr (Table SI 4).

In contrast to the minor response to PFDoA, PFOA induced the largest and broadest change in gene expression in the entire study at 48 hr. For this timepoint, we observed a change in gene expression of a fluoride exporter, central metabolism enzymes, ribosomal proteins, molecular chaperones, autoinducer-2, and motility components induced by PFOA (Figure 8; SI Figure 6; Table SI 3; Table SI 12; Table SI 13, Table SI 14).

**Figure 8.**
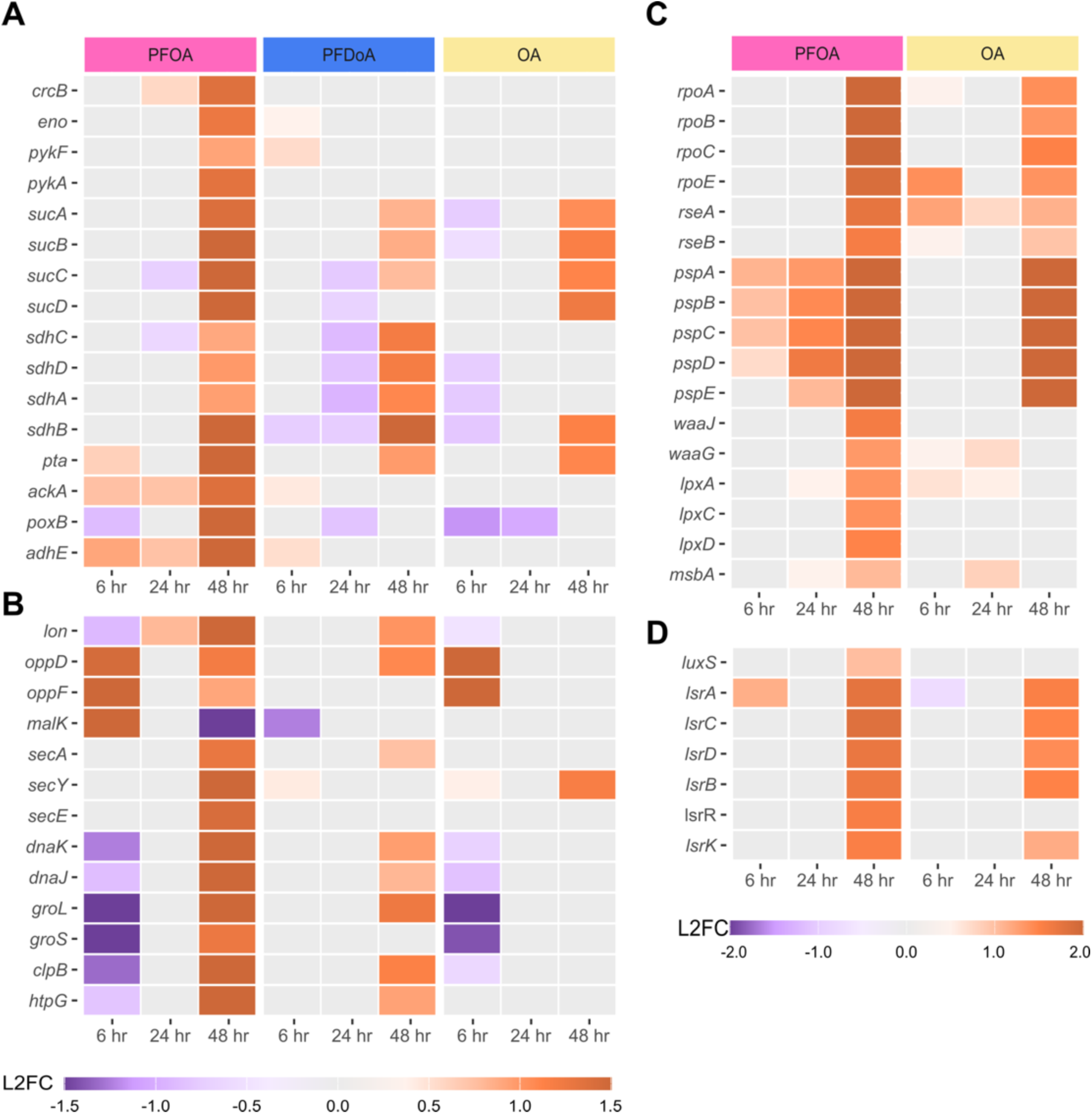
Select genes with differential expression at 48 hr. Heatmaps display differential expression of select genes (padj < 0.05) encoding (A) a fluoride exporter and metabolic enzymes, (B) ATPases and the molecular chaperone network, (C) sigma factors and envelope stress response, and (D) the AI-2 signaling system at select time points in the time course (6, 24, 48 hr). Expression is represented by log_2_ fold change (L2FC) within a gene compared to untreated control in a purple to orange scale displayed by legend color bar below each heatmap. Note heatmaps in A and B share the same L2FC scale and heatmaps in C and D share a separate scale. PFCA or NFCA treatment is represented by color coded bars at the top of the heatmap: PFOA (pink), PFDoA (blue), and OA (yellow). Genes with no significant change in expression are represented by gray boxes.

High concentrations of environmental fluoride can inhibit metabolism, ribosomes, and protein processing, in addition to altering pH and inducing oxidative stress in *E. coli* (45, 78). Similar molecular mechanisms of oxidative stress along with membrane permeability have been demonstrated by PFAS toxicity (38, 95, 96). Exporters, including the selective Fluc family fluoride ion channels, mediate fluoride resistance and protect bacteria against toxicity (97). The *crcB* gene of *E. coli* encodes a Fluc fluoride transport channel to expel fluoride out of the cell (45). Moderate upregulation, log_2_ fold change of 1.4, of *crcB* at 48 hr by PFOA suggests a need to export intracellular fluoride potentially accumulated from PFOA throughout the time course (Figure 5 G; Figure 8 A; Table SI 12). Previous studies have observed the partitioning of PFAS with varying chain length into bacterial membranes and subsequent increase in membrane fluidity (95, 96). In addition, *E. coli* has mechanisms to indiscriminately uptake fluoride through membrane ion channels or porins (45, 98). These results suggest a potential chain-length difference in PFCA uptake and subsequent fluoride accumulation, as *crcB* was not differentially expressed by PFDoA at 48 hr. Additional measurements will be needed to further explore this result.

Key components in various biological processes including glycolysis and the citric acid cycle (TCA) enzymes, ribosomal proteins, ATPases, and molecular chaperones were also upregulated at 48 hr by PFOA (Figure 8 A; Table SI 12). Vital for bacterial growth, genes encoding glycolytic enzymes enolase, *eno*, and pyruvate kinases, *pykF* and *pykA*, were moderately upregulated by PFOA at 48 hr (Figure 8 A; SI Figure 6), while *ppsA*, the phosphoenolpyruvate (PEP) synthetase encoding gene, which catalyzes a reverse reaction from pyruvate to PEP, was not differentially expressed. Enolase and kinases can be inhibited by micromolar amounts of fluoride as the ion can complex to metal ion active sites displacing the electronegative substrate (97, 99). Fluoride inhibition of the activity of these key enzymes in central metabolism can result in the suspension of bacterial growth until fluoride is removed (45, 100). Upregulation of glycolytic enzymes at 48 hr provides further evidence of a specific response to possible fluoride accumulation from PFOA and export with *crcB*. Moreover, additional measurements within and beyond the time course could show if *crcB* mediated export of fluoride alleviates possible inhibition. The *sucABCD* operon, encoding TCA enzymes, was significantly upregulated by PFOA at 48 hr, whereas the upstream *sdhCDAB* operon, encoding succinate dehydrogenase, was moderately upregulated, log_2_ fold change of approximately 1.0 and 1.2, by PFOA at PFDoA, respectively (Figure 8 A; SI Figure 6). Conversion of succinyl-CoA to succinate generates GTP indirectly contributing to energy production (101). Succinyl-CoA also plays a role in peptidoglycan biosynthesis by generating precursor lysine and diaminopimelate (102). Overexpression key metabolic enzymes and the fluoride exporter at 48 hr, suggests *E. coli* could be recovering from possible fluoride inhibition during stationary phase. In connection to partial upregulation of glycolytic and TCA enzymes, genes encoding products of mixed-acid fermentation were also overexpressed in response to PFOA at 48 hr (Figure 8 A; SI Figure 6). Upregulation of genes (*pta, ackA, poxB*, *adhE*) encoding enzymes for pyruvate and acetyl-CoA conversion to acetate, succinate, and ethanol, in addition to activation of upstream enolase further corroborates a possible restart of central metabolism (103). Furthermore, production of mixed-acid fermentation products and TCA intermediates, implies an essential metabolic requirement for survival in nutrient-limited stationary phase exacerbated by PFOA.

Growth in the presence of PFCAs induced the overexpression of ribosomal proteins at 48 hr (Figure 5 H). Forty-two genes encoding 50S and 30S ribosomal protein subunits (104) were upregulated by PFOA at 48 hr, and 17 of which were also upregulated by PFDoA (Table SI 12). Though ribosome biogenesis can be reduced in response to nutrient limitations and stress (105), and fluoride can inhibit protein synthesis (77), the observed increase in gene expression of ribosomal proteins could suggest a requirement for machinery to translate new proteins in response to PFCAs late in stationary phase. These results also suggest an increase in ribosome biogenesis in response to the possible decrease in fluoride concentration by *crcB*-mediated export of fluoride. This observation is corroborated by similar findings in a proteomics study of upregulation of ribosomal proteins in anerobic *E. coli* supplemented with PFOA (38).

Several ATPases, including transporters and molecular chaperones, were also upregulated by PFOA at 48 hr despite no significant change in expression at 24 (Figure 8 B; Table SI 12). ATPases are integral to survival and stress adaptation, and PFCA-induced stress or inhibition by fluoride (78, 106, 107) could lead to disruption of energy metabolism and cellular homeostasis. Upregulation of the ATP-dependent Lon protease (Figure 8 B), with roles in metabolic regulation, protein quality control, and degradation of abundant ribosomal proteins (108, 109), at 48 hr, implies a possible need for the protease to degrade misfolded or abnormal proteins to supply amino acids for protein synthesis during stationary phase (110). Gene expression of ABC transporter ATPases in the oligopeptide transport system, *oppD* and *oppF,* and maltose transport, *malK*, (111) varied throughout the time course. Initially, the ATPases were upregulated at 6 hr, not differentially expressed at 24 hr, and moderately upregulated again at 48 hr by PFOA (Figure 8 B). Likewise, a proteomics study found upregulation of the same oligopeptide-binding proteins in *E. coli* cultivated with PFOA (38). The gene encoding the ATPase SecA and other essential components of the bacterial Sec translocase, *secYG*, (112, 113), were also upregulated at 48 hr (Figure 8 B; Table SI 12). As protein secretion is essential in bacteria, the Sec system provides a form of transmembrane translocation to ferry polypeptides from the cytoplasm across the cytoplasmic membrane into the periplasm (114, 115). Overexpression of these genes suggests possible PFOA-induced intracellular protein accumulation and likely saturation of existing Sec-translocases at 48 hr. Moreover, if fluoride possibly inhibited SecA, upregulation at 48 hr implies an increased abundance of SecA transcripts, similar to other ATPases discussed. Although the gene encoding SecB, the protein secretion chaperone that interacts with the Sec translocase, was not even moderately upregulated with a log_2_ fold change of 0.81, other molecular chaperones may assist in protein processing in response to PFOA.

Bacteria employ molecular chaperones as regulators and facilitators of protein folding, transport, and control to maintain cellular homeostasis, especially during environmental stress. ATP-dependent molecular chaperones DnaK, GroEL, and trigger factor (TF) along with cochaperones DnaJ, GrpE, and GroES coordinate the folding of newly synthesized cytosolic proteins in *E. coli* (116). Genes encoding the chaperone network, *dnaKJ, groLS*, *clpB,* and *htpG*, were highly overexpressed by PFOA at 48 hr (Figure 5 I; Figure 8 B; Table SI 12). Conversely, the genes were initially downregulated by PFOA at 6 hr. Molecular chaperones are overexpressed in response to other environmental stressors such as temperature, heavy metals, limited nutrients, and chemical pollutants (117, 118). Upregulation of molecular chaperones genes during stationary phase suggests stress-induced activation in reaction to the presence of PFOA. This is consistent with existing literature that has found PFAS can destabilize and denature proteins (119, 120). Although PFDoA induced moderate overexpression of select chaperones at 48 hr (Figure 8 B), significant upregulation by PFOA of the chaperone network indicates a chain-length dependent response. Moreover, these results also suggest a specific response to PFCAs as there was no significant change in gene expression induced by either NFCA at 48 hr.

Further analysis revealed PFOA triggered various stress responses as genes encoding sigma factors, components of envelope stress response mechanisms, and motility systems of flagellar assembly and chemotaxis were differentially expressed (Figure 8 C, D; SI Figure 7). RNA polymerase (RNAP) core enzyme subunits, *rpoA and rpoBC*, RNAP sigma factors, *rpoE, rpoH, rpoF/fliA,* and *rpoS,* and RpoE regulators, *rseABC,* were highly upregulated by PFOA and only moderately upregulated by OA at 48 hr (Figure 8 C; Table SI 13). As these genes have regulatory roles in the transcription of genes involved in outer membrane stress response, heat shock, flagellar assembly, and stationary phase gene expression, respectively, overexpression suggests heightened regulation of mechanisms employed to mitigate possible stress of PFCAs and NFCAs (121–123). Similar to RNAP and sigma factors, the phage-shock-protein (Psp) operon, *pspABCD*, was also upregulated by PFOA and OA at 48 hr (Figure 8 C; Table SI 13). In contrast to OA, the magnitude of overexpression of the operon by PFOA increased over the time course. As the Psp system can be initiated by a damaged inner membrane (124) and Sec translocase overload (125), an increase in expression throughout the time course by PFOA suggests use of the regulation mechanism to combat possible envelope stress. Our results show additional evidence of PFOA-induced envelope stress as the lipopolysaccharide (LPS) gene clusters responsible for the synthesis of the core oligosaccharide structure, *rfa*, and the lipid A soluble proteins, *lpxA, lpxC,* and *lpxD*, and the ATP-binding LPS transporter, *msbA,* (126) were moderately upregulated at 48 hr (Figure 8 C; Table SI 13). Overexpression of these genes may suggest the need to synthesize LPS components to maintain the outer membrane, despite the high energetic cost. Previous studies have shown that PFAS partition into lipid bilayers and disrupt bacterial membranes, leading to increased permeability (95, 96, 127). Our results suggest *E. coli* is employing several mechanisms to mitigate envelope stress, likely from PFOA.

To adapt to limited resources in stationary phase and respond to external stressors, bacteria will downregulate high energetic-cost processes like motility to reallocate resources to survival (128, 129). We observed downregulation of genes encoding components involved in flagellar assembly, chemotaxis, and extracellular polysaccharide biosynthesis (EPS) at 48 hr by PFOA (Figure 5 J; Figure 8 D; SI Figure 7; Table SI 14). These results suggest PFOA could be inducing a transition from a sessile state towards possible biofilm formation, therefore reducing motility via downregulation of flagellar assembly, as an indirect effect, in order to conserve energetic resources. Notably, DoA-induced upregulation of the same flagellar assembly and chemotaxis genes during mid-exponential phase (SI Figure 7; Table SI 14) suggests these processes may have been used to respond to DoA and to follow nutrient gradients. Moreover, these genes were not significantly changed by OA or PFDoA at any point in the time course, suggesting a specific response to PFOA at 48 hr. Furthermore, genes encoding autoinducer-2 (AI-2), *luxS*, a quorum-sensing (QS) molecule (130) and the ABC transporter *lsr* operon, used for AI-2 uptake, were significantly upregulated by PFOA at 48 hr (Figure 8 D; Table SI 14). AI-2 controls gene expression of chemotaxis and flagellar assembly and has a possible link to biofilm formation (131). A previous study found PFAS induced the production of EPS, biofilm precursor, in *Rhodococcus* as a stress response (132). Although EPS biosynthesis genes are downregulated at 48 hr, with the exception of the *rfb* operon (Table SI 14), additional measurements after 48 hr could provide insight into possible PFOA-induced biofilm formation.

## CONCLUSIONS

In summary, we performed time course RNA-seq to characterize the transcriptional responses of mid-exponential phase and stationary phase *E. coli* to PFCAs, PFOA and PFDoA. Our results suggest 100 μmol/L of either PFCA or respective NFCAs do not statistically impact the growth of *E. coli* at any point in the time course relative to the untreated control. However, differential expression analysis revealed PFOA and PFDoA did induce statistically significant changes in gene expression of the formate regulon and sulfate assimilation at mid-exponential phase and ferrous iron transport, the molecular chaperone network, and motility processes during stationary phase. The time course enabled us to capture the specific and common responses of *E. coli* to PFCAs and NFCAs across time and helped to characterize responses that were specific to a single growth phase or prolonged over time. For instance, our results indicate a requirement for sulfur-containing compounds, like cysteine, during mid-exponential phase as sulfate assimilation components were overexpressed in response specifically to PFDoA. In contrast, overexpression of ferrous iron transporters and amino acid enzymes by PFOA and PFDoA early in stationary phase suggest a time dependent need for ferrous iron and select amino acids. Although differentially expressed at different phases within the time course, sulfur-containing compounds and iron participate in various redox reactions within the cell which could suggest PFCAs induce adjustments to cellular redox. Interestingly, PFOA induced the largest change in gene expression of this study at 48 hr, whereas PFDoA induced a minor change in expression. These results suggest different modes of interactions between *E. coli* and two PFCAs that only differ in length by four carbons but have different physiochemical properties.

These results not only provide additional biological context of the response of *E. coli* to PFCAs at key phases of growth, but possible novel engineering targets to facilitate the development of future environmental monitoring and mitigation strategies. For example, gene expression of numerous small regulatory RNAs and their targets was only significantly changed by PFCAs and not by NFCAs (Figure 7 C), suggesting a unique response to fluorinated compounds that could be utilized in a whole cell biosensor. Additionally, the moderate upregulation of *crcB*, a putative fluoride exporter downstream of a fluoride riboswitch, by PFOA at 48 hr (Figure 8 A), suggests there may have been some removal of fluorine from the PFOA over the time course, warranting further investigation as a mitigation strategy.

Even though these measurements offer a comprehensive transcriptomic snapshot of the relationship between PFCAs and a model bacterium at key growth phases, further characterization efforts would benefit from additional measurements of PFAS with different molecular structures, concentrations, and complex mixtures to explore the extent to which the responses identified here are conserved and identify novel responses. Furthermore, conducting similar studies on environmental microorganisms and microbial communities could provide insight into conserved responses across genera and species and how contamination by PFAS impacts the environment. As technology continues to advance, improved gene expression measurements at the single cell, population, and community level combined with multiomics level analysis would enable more accurate, impactful understanding of PFAS at all levels of biology. This complete picture could aid in the rapid development of inexpensive and easy-to-deploy living measurement systems for *in situ* environmental monitoring and mitigation strategies.

## MATERIALS AND METHODS

### Bacterial strain

*Escherichia coli* MG1655 ATCC 47076 was obtained from the American Type Culture Collection (ATCC). The strain was maintained on Luria-Bertani (LB) agar plates at 37 °C prior to use.

### PFOA, PFDoA, and NFCA preparation

Two PFCAs, perfluorooctanoic acid (96%, PFOA) and tricosafluorododecanoic acid (95%, perfluorododecanoic acid, PFDoA) were used in this study. See Table SI 15 for additional chemical stock information. PFOA and PFDoA were dissolved in dimethyl sulfoxide (DMSO) to prepare a stock solution of 100 mmol/L. Concentrations in the growth medium for growth experiments were 100 μmol/L and 10 100 μmol/L. The final concentration in the growth medium for sequencing experiments was 100 μmol/L. DMSO was present in the growth medium at a final volume fraction of 0.1%. Dodecanoic acid (98%, DoA) was prepared in the same way as the PFCA stocks. Octanoic acid (99%, OA) samples contained an addition of DMSO to the growth medium. All untreated samples contained an equivalent concentration of DMSO and were devoid of PFCA addition.

### *E. coli* growth in PFCA or NFCA containing medium

*E. coli* MG1655 was struck onto an LB agar (BD Difco) plate from a glycerol stock obtained from ATCC. Overnight cultures for each biological replicate were prepared by inoculating 3 mL of M9 minimal medium (133), without PFCA or NFCA addition, with an individual colony incubated at 37 °C, 250 rpm (Table SI 15). For flask-based growth experiments, PFCA stocks, NFCA stocks, or DMSO were added into M9 prior inoculation with *E. coli* cultures as described above. To prepare sample flasks, 50 mL of M9 in sterile 125 mL baffled flasks was inoculated with overnight culture and adjusted to a starting optical density at 600 nm of 0.01 (SI Figure 1; Table SI 1). Cultures were incubated at 37 °C, 250 rpm for 48 hr. Independent biological triplicate flasks were used for each PFCA, NFCA, and untreated control group. To obtain the optical density of each culture, bacterial growth was monitored in each culture with absorbance measurements at 600 nm in technical triplicate throughout the 48 hr period. A Biotek Epoch2 microplate reader was used for absorbance measurements. A two-way mixed ANOVA of optical density measurements was performed in RStudio running R version 4.2.2 with functions from packages tidyverse (v2.0.0) and rstatix (v0.7.2) (134, 135).

### Total RNA Extraction, cDNA Library preparation, and RNA Sequencing

Cell pellets of *E. coli* from each sample type (PFCA, NFCA, or untreated control) were harvested at 6 hr, 24 hr, and 48 hr and normalized to an OD_600_ of 1.0. Samples were spun down at 5,500xg for 7 min at 4 °C, and supernatant was removed prior to storage at -80 °C until total RNA extraction.

Total RNA was extracted from biological triplicates of each PFCA, NFCA, and untreated control at 6 hr, 24 hr, and 48 hr using RNeasy Mini Kit (Qiagen) according to the manufacturer’s instructions with an on-column DNase I digestion for 15 min using an RNase-free DNase Set (Qiagen). Cells were lysed in 200 μL Tris-EDTA buffer solution, pH 8.0 containing 20 g/L lysozyme (Sigma Aldrich) at room temperature for 10 min with vortexing every 2 min. RNA was eluted in 35 μL of molecular grade water (Invitrogen) and stored at -80 °C. The concentration and purity of total RNA were quantified on a Qubit 4 fluorometer (Invitrogen) using Qubit RNA Broad Range Assay Kit, DeNovix D-11 Series Spectrophotometer, and 4200 TapeStation System (Agilent) using RNA ScreenTape Analysis.

Total RNA samples were sent to Azenta Life Sciences (USA) for cDNA library preparation using TruSeq Stranded mRNA kit (Illumina) and next-generation sequencing (Strand-Specific RNA-Seq Service) using an Illumina HiSeq platform for 2x150 bp paired-end reads. Average sequencing depth obtained was 7.81 million. Additional information on sequencing is provided in (Table SI 2).

### Bioinformatic analysis

Bioinformatic analysis was performed on an Amazon Web Service EC2 instance and locally on RStudio. TrimGalore (v0.6.7) with Cutadapt (v3.5) and fastQC (v0.11.9) was used to remove adapter sequences and low-quality reads from each raw read file (136–138). HISAT2 (v2.2.1) (139) was used to align paired-end reads to *E. coli* MG1655 reference genome (GenBank accession GCA_000005845.2). Alignment scores and quality metric scores are shown in Table SI 2. Samtools (v1.13) was used for file conversion of SAM files to sorted BAM files before transcript assembly was performed with Stringtie (v2.2.1) (140, 141). In RStudio running R version 4.2.2, tximport (v1.26.1) prepared count tables from Stringtie transcript-level estimates (134, 135, 142). DESeq2 (v1.38.3) was used to perform differential gene expression analysis with a negative binomial distribution model (49). The time specific untreated control served as the control baseline for all pairwise comparisons. In this study, we consider genes to be differentially expressed if the Benjamini-Hochberg adjusted p-value (padj) is less than 0.05. We mark moderate and significant differential expression with absolute values of the log_2_ fold change greater than or equal to 1.2 and 1.5, respectively. Gene ontology and Kyoto Encyclopedia of Genes and Genomes pathway over-representation analyses were performed using clusterProfiler (v4.6.2) using the *E. coli* K12 organism package, org.EcK12.eg.db, (v3.19.1) and KEGG organism eco (143, 144). Additional data curation was performed using EcoCyc (145).

## DATA AVAILABILITY

The data presented in this study have been deposited in the National Center for Biotechnology Information’s Gene Expression Omnibus (146) and are accessible through GEO Series accession number GSE288896 (https://www.ncbi.nlm.nih.gov/geo/query/acc.cgi?acc=GSE288896).

## Author contributions

M.E.W. conceived the project, designed experiments, conducted data analysis, and wrote the first draft with input from S.W.S. O.B.V. designed growth experiments. M.E.W. and S.W.S edited the manuscript with input from all authors.

The authors would like to thank Elizabeth Strychalski, Allison Yaguchi, Justin Wagner, Svetlana Ikonomova, and Zoila Jurado Quiroga for insightful discussions throughout the project and on the manuscript.

Figure 1, Figure 5, SI Figure 1, and SI Figure 6 were created with BioRender.com.

## Disclaimer

Certain commercial entities, equipment, or materials may be identified in this document to describe an experimental procedure or concept adequately. Such identification is not intended to imply recommendation or endorsement by the National Institute of Standards and Technology, nor is it intended to imply that the entities, materials, or equipment are necessarily the best available for the purpose. Official contribution of the National Institute of Standards and Technology; not subject to copyright in the United States.

