## Supplemental figures 1-7 for "Comparative Transcriptomic Analysis of Perfluoroalkyl Substances-Induced Responses of Exponential and Stationary Phase *Escherichia coli*"

<u>List of SI Figures</u>


1. SI Figure 1. Visual overview of culturing and sampling procedure in biological triplicate.

2. SI Figure 2. Growth of *E. coli* MG1655 in M9 at 37 °C in the presence of 10 µmol/L of PFOA (pink), PFDoA, (blue), OA (yellow), or DoA (green) displayed as $OD_{600}$ over time (hr).

3. SI Figure 3. Principal component analysis of transcriptome samples displays sample variation of treated and untreated samples at 6 hr (square), 24 hr (circle), and 48 hr (triangle).

4. SI Figure 4. Transcriptomic profiles of *E. coli* grown with NFCAs.

5. SI Figure 5. Differential gene expression comparison between PFCAs and NFCAs by chain length over time.

6. SI Figure 6. Schematic of metabolic pathways differentially expressed in stationary phase *E. coli* in response to PFCAs.

7. SI Figure 7. Select motility genes with differential expression over the time course.

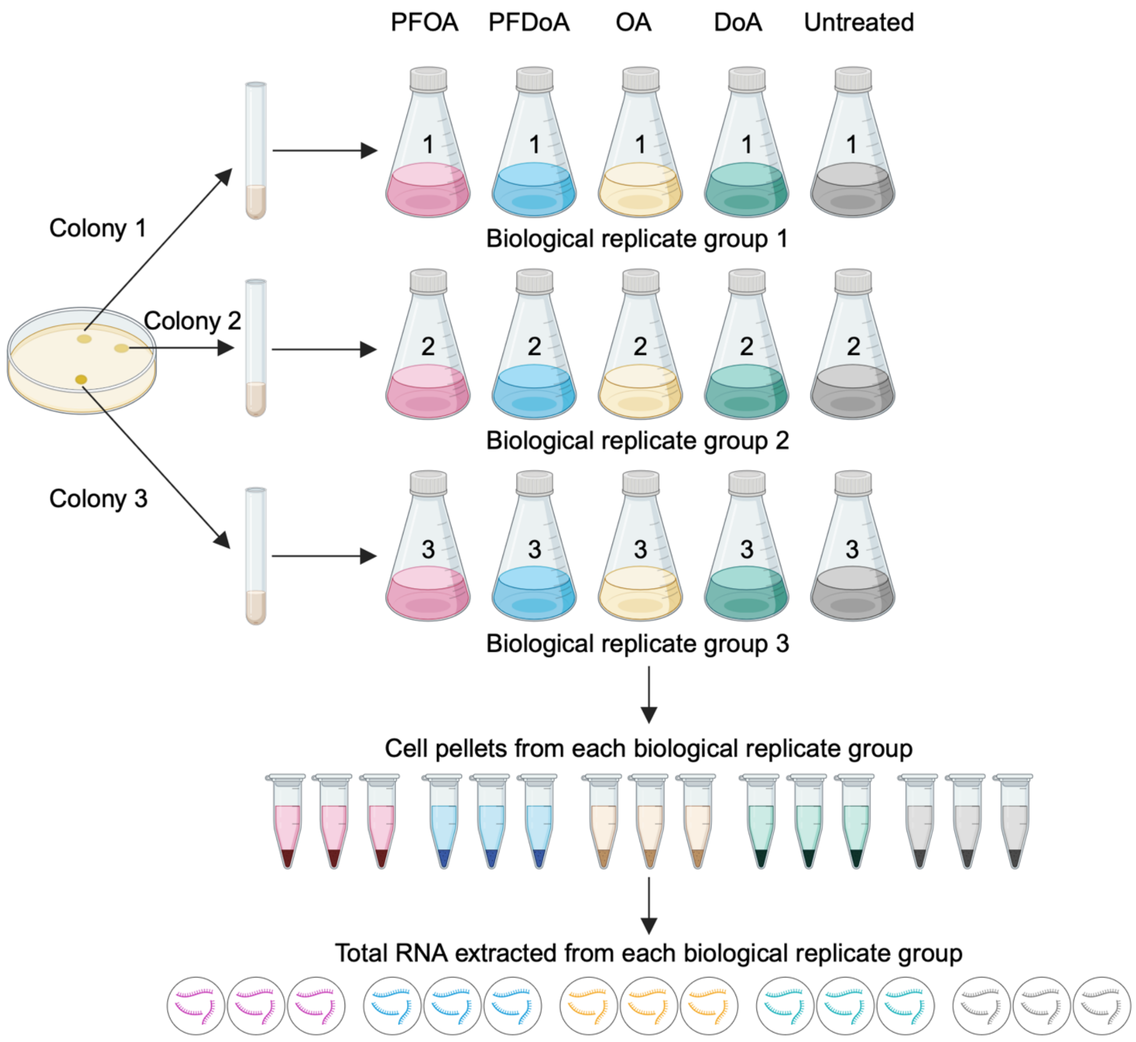


**SI Figure 1.** Visual overview of culturing and sampling procedure in biological triplicate with each experimental condition of perfluorooctanoic acid (PFOA, pink), perfluorododecanoic acid (PFDoA, blue), octanoic acid (OA, yellow), dodecanoic acid (DoA, green), and untreated (gray).

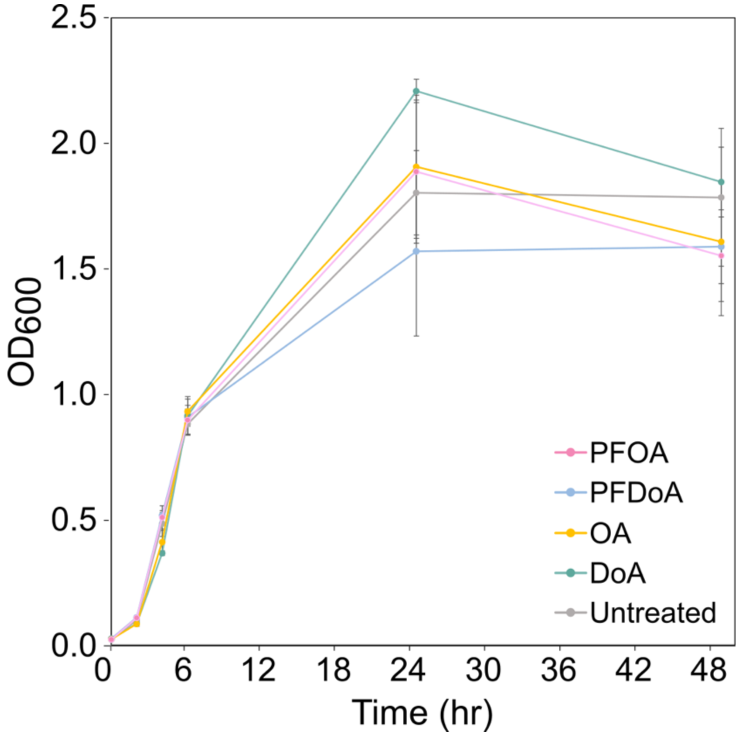


**SI Figure 2.** Growth of *E. coli* MG1655 in M9 at 37 °C in the presence of 10 μmol/L of PFOA (pink), PFDoA, (blue), OA (yellow), or DoA (green) displayed as $OD_{600}$ over time (hr). Untreated cells containing an equivalent concentration of DMSO are displayed in gray. Error bars represent standard deviation of biological triplicates.

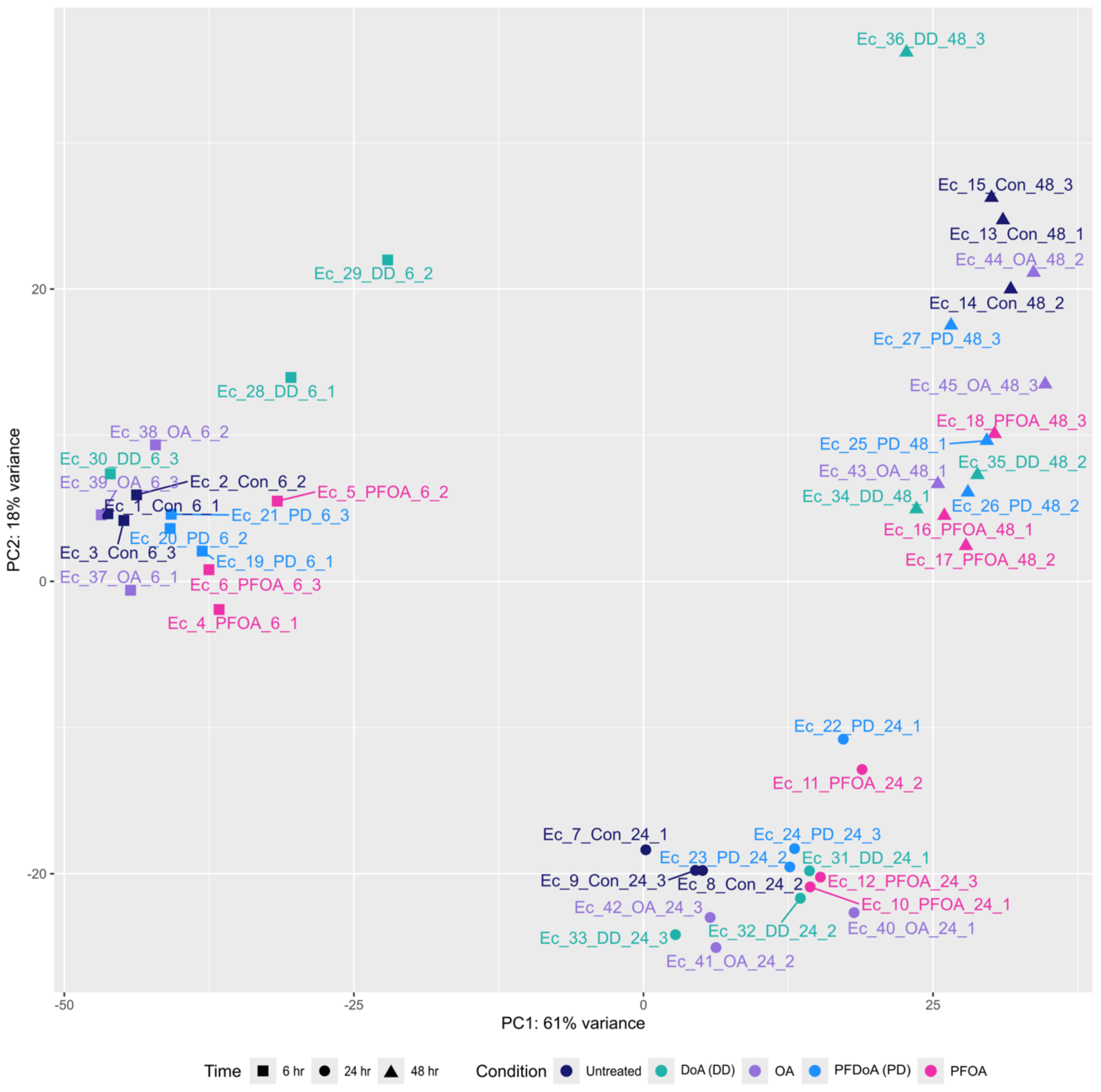


**SI Figure 3. Principal component analysis of transcriptome samples displays sample variation of treated and untreated samples at 6 hr (square), 24 hr (circle), and 48 hr (triangle).** Sample type is represented by color: PFOA (pink), PFDoA denoted as PD (blue), OA (purple), DoA denoted as DD (green), and untreated control (navy). Sample table with identifiers in SI Table 2.

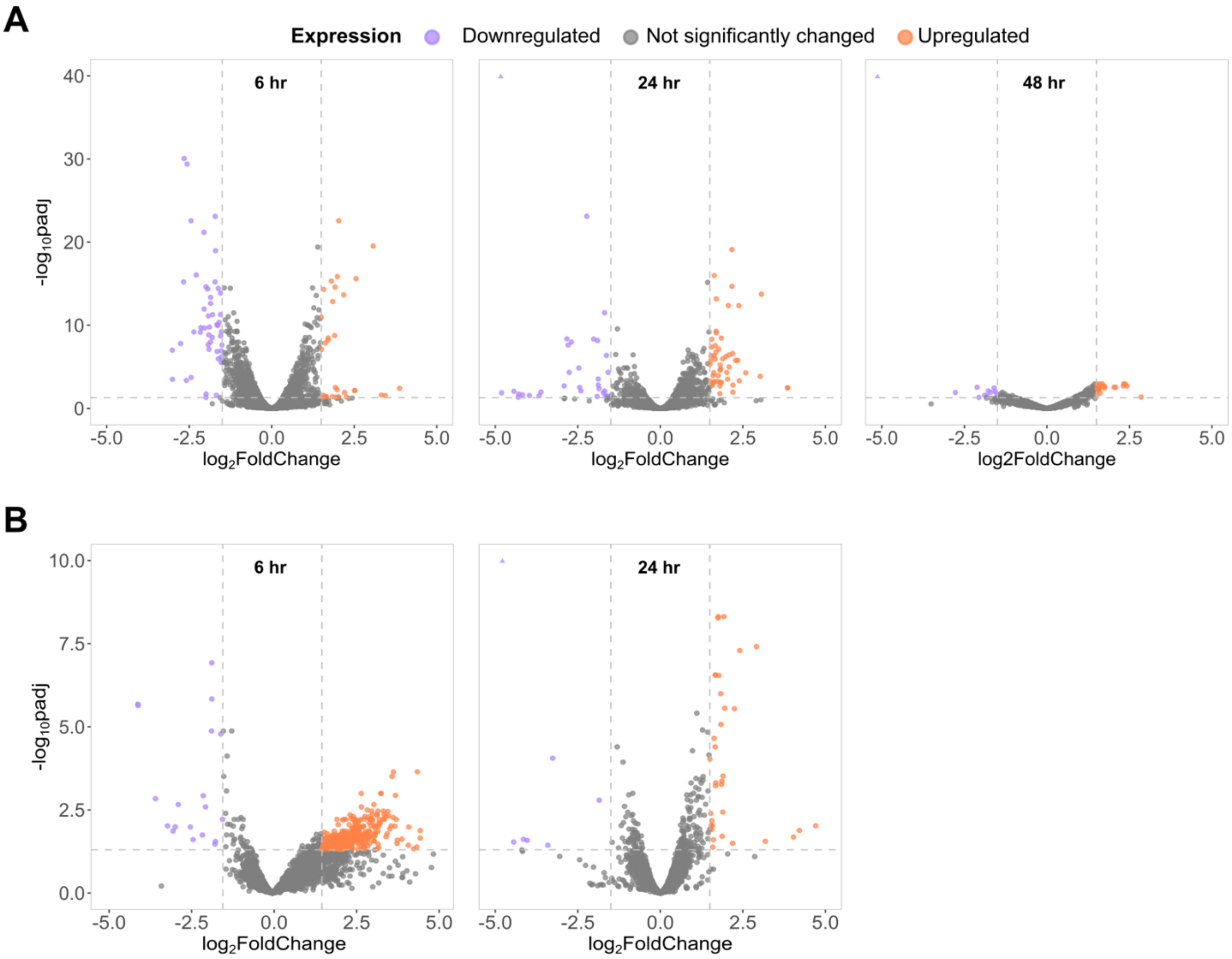


**SI Figure 4. Transcriptomic profiles of *E. coli* grown with NFCAs.** Volcano plots show the global transcriptional response of *E. coli* to (A) OA and (B) DoA in comparison to an untreated control group at 6 hr, 24 hr, and 48 hr. Dashed lines represent a significance cutoff for differential expression of an adjusted p-value (padj) of less than 0.05 and absolute value of the $\log_2$ foldchange greater than 1.5. Upregulated and downregulated genes are denoted with purple and orange, respectively, and genes not significantly changed by NFCA treatment are gray. Differentially expressed genes with $\log_2$ foldchange and or padj values outside of the plot limits are represented by orange or purple triangles in the upper corners of each plot if applicable.

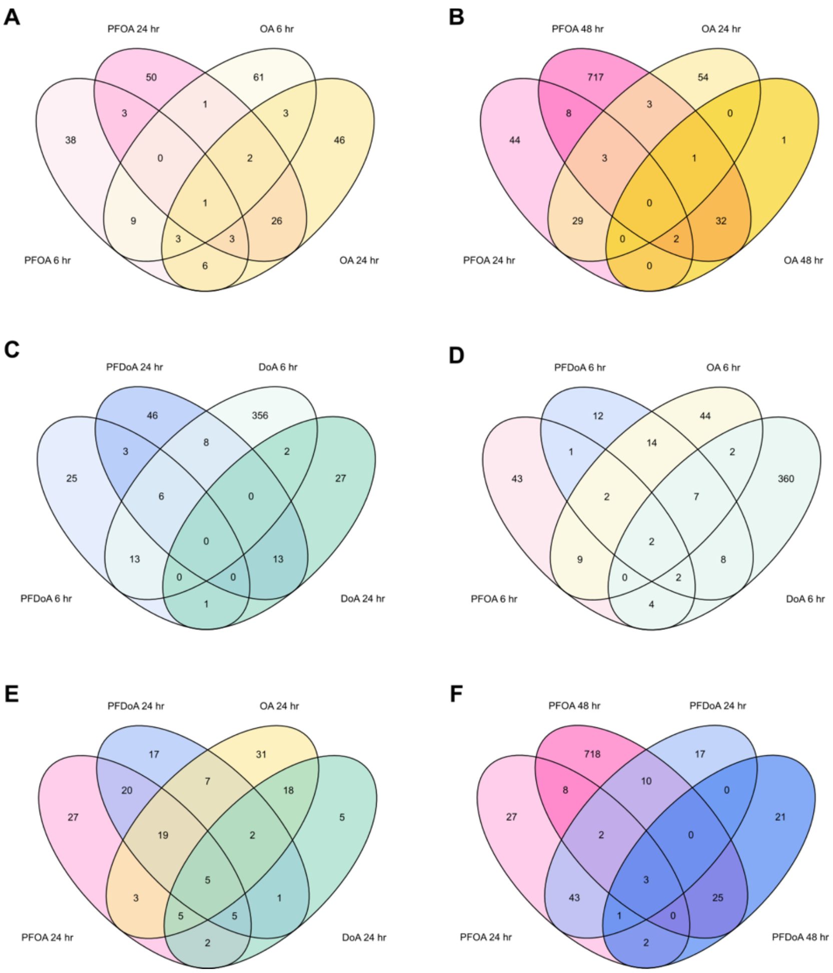


**SI Figure 5**. **Differential gene expression comparison between PFCAs and NFCAs by chain length over time.** Venn diagrams display the number of differentially expressed genes uniquely or commonly induced in *E. coli* by the labeled treatment. Unions display genes differentially expressed by both treatments at more than one time point. Genes in the outer circles are only differentially expressed at a single time point by a single treatment. (A) Comparison of differentially expressed genes by PFOA or OA at 6 hr and 24 hr. (B) Comparison of differentially expressed genes by PFOA or OA at 24 hr and 48 hr. (C) Comparison of differentially expressed genes by PFDoA or DoA at 6 hr and 24 hr.(D) Comparison of differentially expressed genes by PFOA, PFDoA, OA, or DoA at 6 hr. (E) Comparison of differentially expressed genes by PFOA, PFDoA, OA, or DoA at 24 hr. (F) Comparison of differentially expressed genes by PFOA or PFDoA at 24 hr and 48 hr.

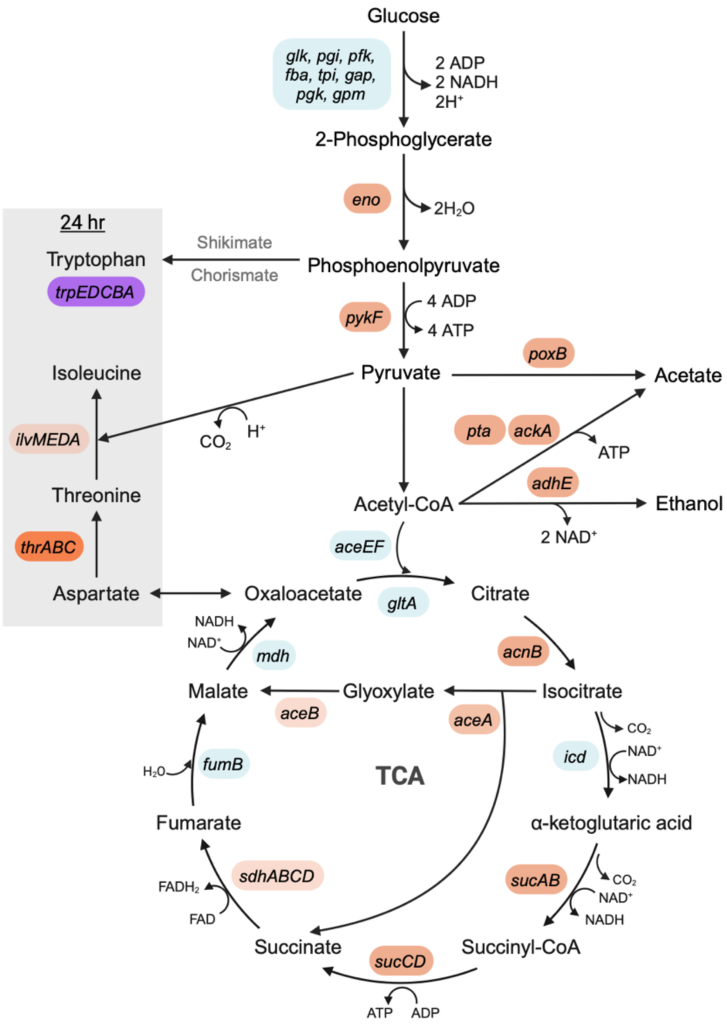


**SI Figure 6. Schematic of metabolic pathways differentially expressed in stationary phase *E. coli* in response to PFCAs.** The reactions and synthesis pathways of amino acids including aspartate, threonine, isoleucine, and tryptophan differentially expressed at 24 hr by PFOA and PFDoA are highlighted with a gray background. The pathways differentially expressed at 48 hr by PFOA include glycolysis, the citric acid cycle (TCA), and fermentation reactions to acetate and ethanol. Differentially expressed genes (padj < 0.05) are shaded in orange (upregulated) or purple (downregulated) with corresponding $log_2$ foldchanges displayed in Figure 7, Figure 8, Table SI 8, and Table SI 9. Genes with no significant change in expression (padj > 0.05 and or |log2 foldchange| < 1.2) are shaded in blue.

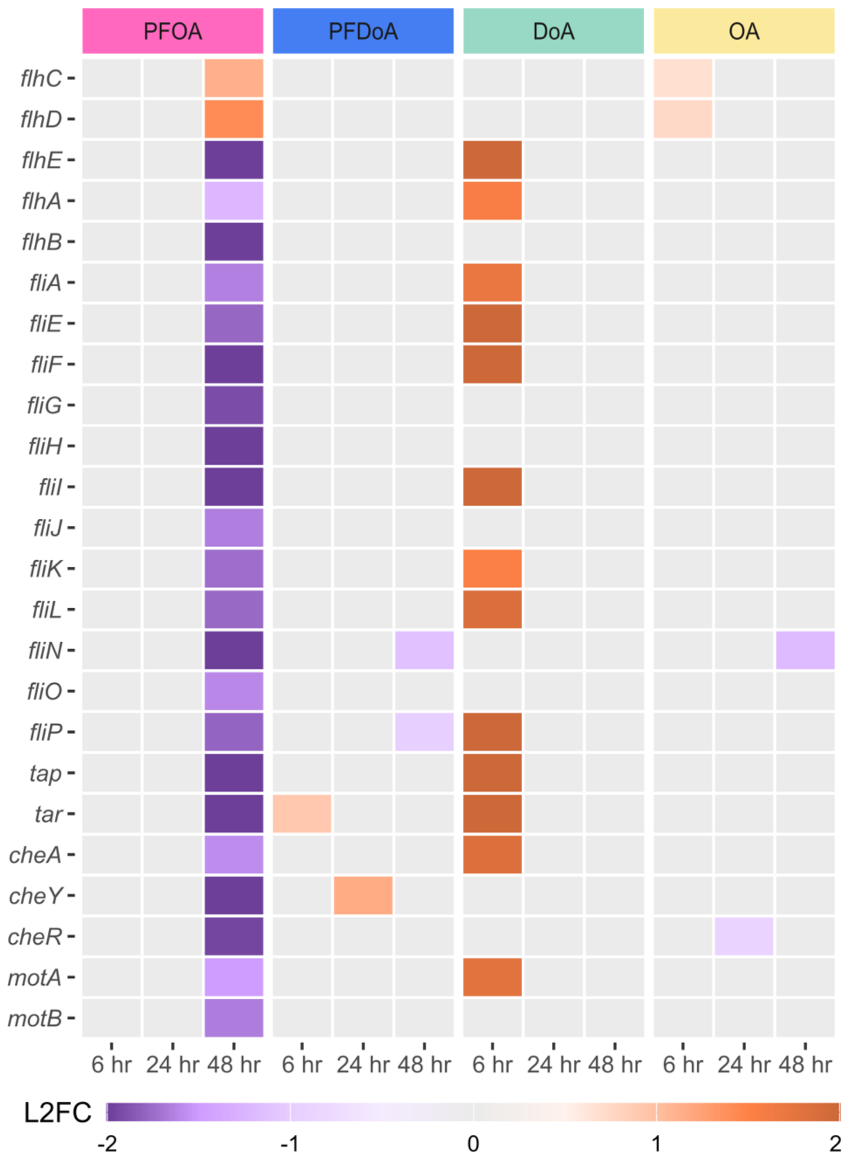


**SI Figure 7. Select motility genes with differential expression over the time course.** Heatmaps display differential expression of select genes (padj < 0.05) encoding motility processes including flagellar assembly and chemotaxis at select time points in the time course (6, 24, 48 hr). Expression is represented by $log_2$ fold change (L2FC) within a gene compared to untreated control in a purple to orange scale displayed by legend color bar below each heatmap. PFCA or NFCA treatment is represented by color coded bars at the top of the heatmap: PFOA (pink), PFDoA (blue), DoA (green), and OA (yellow). Genes with no significant change in expression are represented by gray boxes.
